# Genetic ablation of neuronal mitochondrial calcium uptake impedes Alzheimer’s disease progression

**DOI:** 10.1101/2023.10.11.561889

**Authors:** Pooja Jadiya, Elena Berezhnaya, Devin W. Kolmetzky, Dhanendra Tomar, Henry M. Cohen, Shatakshi Shukla, Manfred Thomas, Salman Khaledi, Joanne F. Garbincius, Liam Kennedy, Oniel Salik, Alycia N. Hildebrand, John W. Elrod

## Abstract

Loss of _m_Ca^2+^ efflux capacity contributes to the pathogenesis and progression of Alzheimer’s disease (AD) by promoting mitochondrial Ca^2+^ (_m_Ca^2+^) overload. Here, we utilized loss-of-function genetic mouse models to causally evaluate the role of _m_Ca^2+^ uptake by conditionally deleting the mitochondrial calcium uniporter channel (mtCU) in a robust mouse model of AD. Loss of neuronal _m_Ca^2+^ uptake reduced Aβ and tau-pathology, synaptic dysfunction, and cognitive decline in 3xTg-AD mice. Knockdown of *Mcu* in an *in vitro* model of AD significantly reduced matrix Ca^2+^ content, redox imbalance, and mitochondrial dysfunction. The preservation of mitochondrial function rescued the AD-dependent decline in autophagic capacity and protected neurons against amyloidosis and cell death. This was corroborated by *in vivo* data showing improved mitochondrial structure and apposition in AD mice with loss of neuronal *Mcu*. These results suggest that inhibition of neuronal _m_Ca^2+^ uptake represents a powerful therapeutic target to impede AD progression.

## Introduction

The mechanisms underlying Alzheimer’s disease (AD) pathogenesis remain at-large and this gap in knowledge is a significant barrier to the development of effective therapeutic interventions. AD is associated with mitochondrial dysfunction (Du *et al*, 2008; Gillardon *et al*, 2007; Rossi *et al*, 2020) and altered neuronal intracellular calcium (_i_Ca^2+^) homeostasis (Begley *et al*, 1999; Mattson *et al*, 1992). Many studies point to increased [_i_Ca^2+^] as an upstream mechanism that may promote β-amyloid (Aβ) and tau pathology (Barrett *et al*, 2014; Kipanyula *et al*, 2012; Verma *et al*, 2017). Excessive _i_Ca^2+^ can result from the dysregulation of Ca^2+^ channels, such as calcium homeostasis modulator 1 (CALHM1) (Dreses-Werringloer *et al*, 2008), amino-3-hydroxy-5-methyl-4-isoxazolepropionic acid receptors (AMPAR) (Chang *et al*, 2006), or N-methyl-D-aspartate receptors (NMDAR) (Mattson *et al*., 1992). Further, alterations in store-operated calcium entry (SOCE) (Tong *et al*, 2016) and increased ryanodine receptor (RyR) (Paula-Lima *et al*, 2011) and inositol trisphosphate receptor (IP3R) activity are reported to increase ER Ca^2+^ content in the context of AD (Ferreira *et al*, 2015). This coupled with reports of increased ER-mitochondria coupling (Area-Gomez *et al*, 2018; Area-Gomez *et al*, 2012) suggests that mitochondrial calcium (_m_Ca^2+^) overload may be a critical event in AD progression.

In contrast, other studies suggest that Aβ pathologies are upstream of impaired neuronal _i_Ca^2+^ handling (Calvo-Rodriguez *et al*, 2019) and mitochondrial dysfunction (Lustbader *et al*, 2004), leading to cognitive impairment, synaptic dysfunction, and neurodegeneration in AD. For example, mutations in PS1 are reported to cause RyR hyperactivity (Lacampagne *et al*, 2017) and increased IP3R channel gating (Cheung *et al*, 2010; Cheung *et al*, 2008) which leads to elevated _i_Ca^2+^ signaling. The familial AD (FAD) PS1^M146L^ mutant interacts with the IP3R, increasing Ca^2+^ release from the ER, stimulating APP processing (Cheung *et al*., 2008), and contributing to neuronal pathology. Studies suggest that increased _i_Ca^2+^ signaling through the IP3R-PS1 interaction is a disease-specific mechanism in FAD (Cheung *et al*., 2010), enhancing the production of reactive oxygen species and contributing to AD pathogenesis (Muller *et al*, 2011). Similarly, post-translational remodeling (PKA phosphorylation, oxidation, and nitrosylation) of neuronal RyR2 channels in human AD patients and murine models of AD induces ER-Ca^2+^ leak resulting in calpain activation, tau phosphorylation, and cognitive deficits (Lacampagne *et al*., 2017). Thus, the literature supports the rationale that calcium dysregulation can both precede and follow Aβ and tau pathology.

Elevations in _i_Ca^2+^ are theorized to be rapidly integrated into mitochondria due to the high electromotive force across the inner mitochondrial membrane (IMM) generated by the electron transport chain (Δψ = ∼ -160mv). Given the driving force for _m_Ca^2+^ entry and the highly dynamic nature of [_i_Ca^2+^], neuronal mitochondria require a tightly regulated _m_Ca^2+^ exchange system (Billups & Forsythe, 2002; Cardenas *et al*, 2010). Ca^2+^ enters the mitochondrial matrix via the mitochondrial calcium uniporter channel complex (mtCU) and modulates key metabolic control points in the TCA cycle (Denton *et al*, 1972; Denton *et al*, 1978). It is postulated that _m_Ca^2+^ is the signal that matches mitochondrial energy production to neuronal energy demand. Excessive [_m_Ca^2+^] causes increased oxidative stress, mitochondrial dysfunction, opening of the mitochondrial permeability transition pore (mPTP), and ultimately neuronal death (Du *et al*., 2008; Luongo *et al*, 2015; Nakagawa *et al*, 2000; Szalai *et al*, 1999). We recently reported that _m_Ca^2+^ overload due to decreased expression of NCLX, the mitochondrial sodium-calcium exchanger and primary mechanism for _m_Ca^2+^ efflux, contributes to AD progression in the 3xTg-AD mouse model and rescue of NCLX expression was sufficient to abrogate behavioral and histopathological hallmarks of AD (Jadiya *et al*, 2019). We also noted proteomic remodeling of the mtCU in sporadic AD patients (Jadiya *et al*., 2019). In support of our findings, a recent study showed increased neuronal _m_Ca^2+^ levels correlated with plaque deposition in the APP/PS1-Tg mouse model and other reports have implicated _m_Ca^2+^ overload in the activation of cell death and neurodegeneration (Granatiero *et al*, 2019; Jadiya *et al*., 2019; Kostic *et al*, 2015; Logan *et al*, 2014; Luongo *et al*., 2015; Qiu *et al*, 2013). For example, deletion of cyclophilin D (CypD), a necessary activator of the mPTP, has been shown to prevent mitochondrial dysfunction and memory impairments in AD models (Du *et al*., 2008). Further, a recent study utilizing a *C. elegans* AD model, *sel-12* (homolog of presenilin) mutant, observed increased ER-mitochondria Ca^2+^ signaling resulting in increased ROS (reactive oxygen species) generation and neuronal dysfunction (Sarasija *et al*, 2018). Similarly, an AD-linked presenilin mutation (PS1) was reported to increase inositol triphosphate (IP3)-mediated Ca^2+^ transients (Stutzmann *et al*, 2004) and ER-mitochondrial contacts in AD (Area-Gomez *et al*., 2012), both of which could contribute to _m_Ca^2+^ overload. These studies strengthen the notion that _m_Ca^2+^ overload is central to AD progression.

*Mitochondrial calcium uniporter* (*MCU*) encodes the pore-forming component of the mtCU complex and is required for channel function (Baughman *et al*, 2011; De Stefani *et al*, 2011). Fibroblasts isolated from AD patients display a significant increase in MCU protein expression compared to controls (Perez *et al*, 2018). Notably, overexpression of MCU by stereotaxic injection of adenovirus into the cortex of mice resulted in increased mitochondrial Ca^2+^ levels, neuronal cell death, and gliosis (Granatiero *et al*., 2019), suggesting that _m_Ca^2+^ overload alone is sufficient to promote brain pathology. To define how changes in _m_Ca^2+^ uptake causally contribute to the development of AD, we generated 3xTg-AD mutant mice with neuronal-specific deletion of *Mcu* (*Mcu^fl/fl^* x Camk2a-Cre x 3xTg-AD). Loss of acute _m_Ca^2+^ uptake prevents cognitive decline and ameliorates proteotoxic hallmarks of disease including Aβ aggregation and tau phosphorylation in 3xTg-AD mice. In an *in vitro* model of AD we show that inhibition of _m_Ca^2+^ uptake ameliorates mitochondrial dysfunction and ROS generation rescuing autophagy impairments and protects neurons against amyloidosis and cell death. These data demonstrate that targeting _m_Ca^2+^ uptake is a novel therapeutic to impede AD development and progression.

## Results

### Loss of neuronal _m_Ca^2+^ uptake improves AD-associated cognitive deficits

To examine the role of _m_Ca^2+^ uptake in AD pathology, we generated a neuronal-specific knockout of *Mcu* in the 3xTg-AD mutant mouse model. 3xTg-AD mice develop robust disease including both Aβ and tau pathology and memory decline. 3xTg-AD mice were crossed to mice expressing neuron-restricted Cre recombinase (Tsien *et al*, 1996) harboring the *Mcu* floxed allele (*Mcu*^fl/fl^ x Camk2a-Cre) (Luongo *et al*., 2015) to delete *Mcu* specifically from frontal cortex and hippocampal neurons in AD mice (3xTg-AD x *Mcu*^fl/fl^ x Camk2a-Cre, **Figure 1A**). Loss of MCU expression from the frontal cortex of animals harboring both *Mcu^fl/fl^* and Camk2a-Cre alleles was confirmed at both 2 and 12 months via Western blot, validating the experimental model system (**Figure 1B-C, Figure S1 O-P**). No significant changes were observed in the expression of other mitochondrial calcium uniporter channel complex (mtCU)-associated components in the 3xTg-AD x *Mcu*^fl/fl^ x Camk2a-Cre mice compared to 3xTg-AD x Camk2a-Cre mice. However, in 12-month-old mice harboring the 3xTg-AD alleles there was a reduction in MCUB, MICU1, MICU3, and NCLX expression when compared to Camk2a-Cre controls (**Figure 1B-C, S1 A-N**). This decrease in the expression of mtCU regulators is likely related to elevated _m_Ca^2+^ levels, which are observed during AD progression and validates reported results in non-familial AD patient samples (Calvo-Rodriguez *et al*, 2020; Jadiya *et al*, 2023a; Jadiya *et al*., 2019; Jadiya *et al*, 2023b). Previous studies have shown that homozygous MICU1 deletion in neurons leads to altered Ca^2+^ homeostasis and progressive motor and cognitive dysfunction (Singh *et al*, 2022). As MICU1 and MICU3 regulate MCU-dependent Ca^2+^ uptake (Ashrafi *et al*, 2020; Singh *et al*., 2022), their downregulation in the transgenic AD model suggests that MICU1 and MICU3 deficiency may promote aberrant _m_Ca^2+^ uptake. Next, we examined _m_Ca^2+^ uptake and calcium retention capacity (CRC) using mitochondria isolated from the frontal cortex of 12-month-old mice with MCU deletion in the 3xTg-AD background. 3xTg-AD mice demonstrated reduced CRC with no significant increase in _m_Ca^2+^ uptake rate relative to controls (**Figure 1D-F**). Loss of neuronal MCU caused near complete loss of _m_Ca^2+^ uptake and CRC (**Figure 1D-F**).

**Figure 1.**
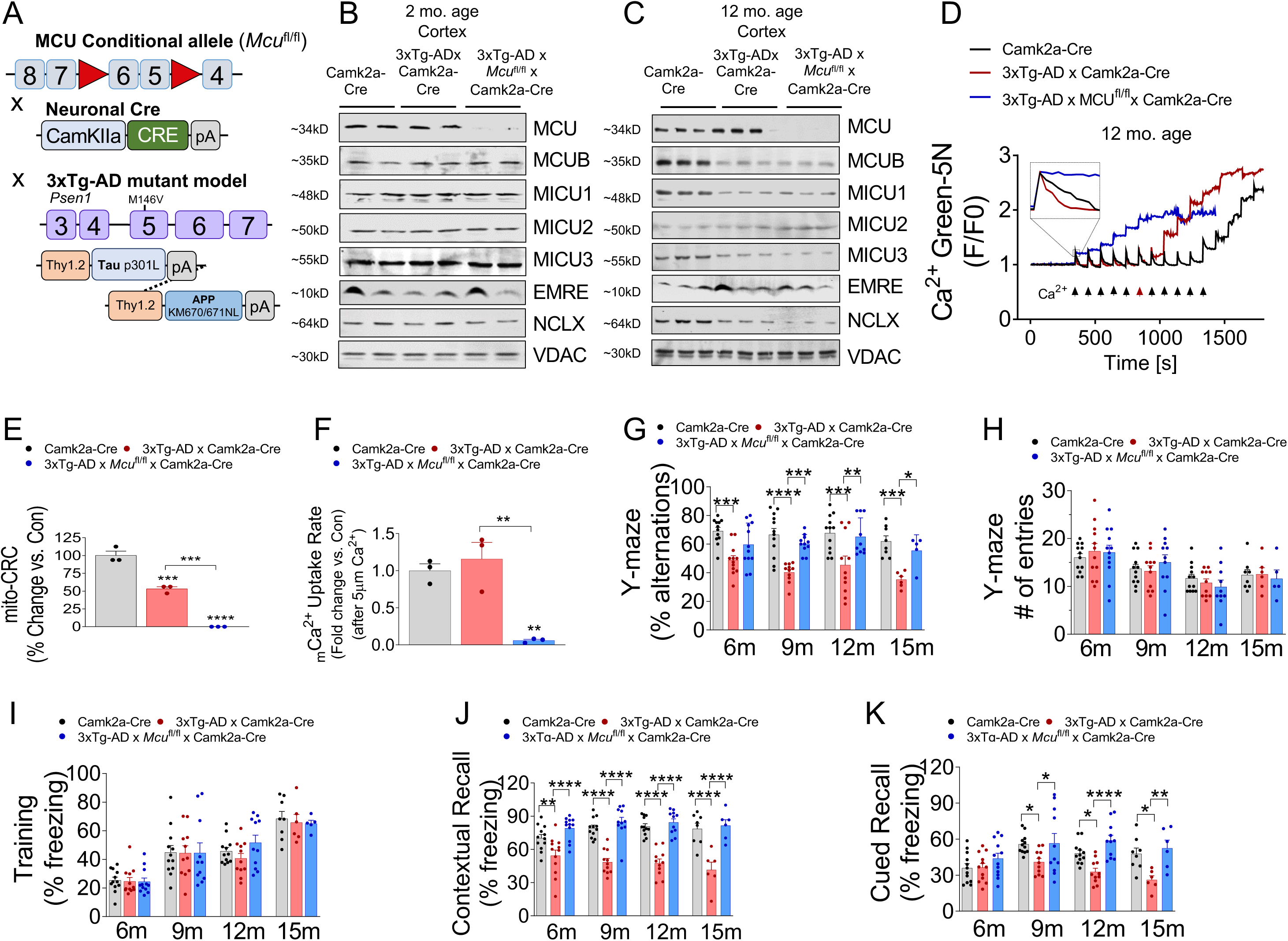
Loss of neuronal _m_Ca^2+^ uptake improves AD-associated cognitive deficits. **A** Schematic of *Mcu* knockout 3xTg-AD mutant mouse gene-targeting strategy. **B** Western blots for MCU expression and proteins associated with _m_Ca^2+^ exchange in tissue isolated from the cortex of 2 months old 3xTg-AD x *Mcu*^fl/fl^ x Camk2a-Cre mice compared to age-matched control. MCU, mitochondrial calcium uniporter; MCUB, mitochondrial calcium uniporter β subunit; MICU1, mitochondrial calcium uptake 1; MICU2, mitochondrial calcium uptake 2; MICU3, mitochondrial calcium uptake 3; EMRE, Essential MCU Regulator; NCLX, Mitochondrial Na+/Ca2+ Exchanger. VDAC, Voltage-dependent anion channel served as mitochondrial loading controls. **C** Western blots for MCU expression and proteins associated with _m_Ca^2+^ exchange in tissue isolated from the cortex of 12 months old 3xTg-AD x *Mcu*^fl/fl^ x Camk2a-Cre mutant mice compared to age-matched control. **D** Representative traces for _m_Ca^2+^ retention capacity (CRC). **E** Percent change in CRC of 3xTg-AD × Camk2a-Cre and 3xTg-AD × *Mcu*^fl/fl^ × Camk2a-Cre vs. Camk2a-Cre control. **F** _m_Ca^2+^ uptake rate expressed as fold-change vs. Camk2a-Cre con. calculated after first 5-µm Ca^2+^ bolus. **G, H** Y-maze spontaneous alternation test. **G** Percentage of spontaneous alternation. **H** Total number of arm entries. **I–K** Fear-conditioning test. **I** Freezing responses in the training phase. **J** Contextual recall freezing responses, **K** Cued recall freezing responses. n = individual dots shown for each group in all graphs. All data presented as mean ± SEM; ****p < 0.0001, ***p < 0.001, **p < 0.01, *p < 0.05; one-way ANOVA with Sidak’s multiple comparisons test.

Next, we sought to assess whether loss of neuronal _m_Ca^2+^ uptake was sufficient to reduce or prevent cognitive decline in 3xTg-AD mice. Mice from all groups (Camk2a-Cre, 3xTg-AD x Camk2a-Cre, and 3xTg-AD x *Mcu*^fl/fl^ x Camk2a-Cre) were tested for spatial memory by Y-maze at 6, 9, 12 and 15 months of age. Loss of neuronal MCU significantly improved AD-associated impairments in spatial working memory evidenced by increased alternations from 9-15 months of age (**Figure 1G**). Importantly, we did not see any significant differences in locomotor function between groups, as indicated by an equivalent number of arm entries (**Figure 1H**).

Next, we used a fear-conditioning paradigm to assess changes in contextual and cued fear conditioning. Freezing behavior was measured in response to either the training environment (contextual) or a tone previously paired with foot shock (cued). No differences in freezing response were observed during the training session between groups throughout the study, demonstrating no measurable effect on locomotor activity or differences in baseline behavior (**Figure 1I**). 3xTg-AD mice displayed impairments in contextual and cued recall from 6-15 months and 12-15 months of age, respectively (**Figure 1J and 1K**). AD mice with neuronal deletion of *Mcu* exhibited a significant improvement in contextual and cued recall from 6-15 months of age (**Figure 1J and 1K**). Importantly, loss of neuronal MCU alone (*Mcu*^fl/fl^ x Camk2a-Cre) did not result in cognitive impairment in Y-maze testing (**Figure S1 Q-R**) and demonstrated a slight decrease in contextual recall at 12 months of age (**Figure S1 S-U**), perhaps suggesting a physiological role for _m_Ca^2+^ uptake in specific neuronal populations involved in long-term amygdala and/or hippocampus memory in aged mice. (Curzon *et al*, 2009).

### Genetic ablation of neuronal _m_Ca^2+^ uptake attenuates Aβ accumulation in 3xTg-AD mice

We examined whether ablating neuronal _m_Ca^2+^ uptake is protective against amyloid pathology. Cortical and hippocampal homogenates isolated from aged mice (15 months) were assayed by ELISA for Aβ_1-40_ and Aβ_1-42_ peptide levels in both RIPA-soluble and -insoluble fractions. Loss of neuronal _m_Ca^2+^ uptake in AD mice resulted in a significant decrease in both soluble and insoluble Aβ_1-40_ and Aβ_1-42_ levels in cortex and hippocampus, without any change in the Aβ_42/40_ ratio (**Figure 2A-D**; **S2A-B**). Similarly, loss of neuronal _m_Ca^2+^ uptake caused a significant reduction (∼40%) in amyloid deposits in AD mice (**Figure 2E, F**). To further investigate the mechanism by which genetic loss of *Mcu* reduces Aβ burden, we examined the expression of all secretases involved in APP processing including α, β and γ-secretase (**Figure 2G; S2C)**. As expected, aged 3xTg-AD mice demonstrated increased expression of total APP, β (BACE-1) and γ-secretase complex components (PS1, NCT and APH1 subunit), whereas α-secretase (ADAM10) expression was reduced. Interestingly, deletion of neuronal *Mcu* from 3xTg-AD mice reverted the increased expression of γ-secretase complex associated proteins presenilin-1 (PS-1) and Nicastrin (NCT) (**Figure 2G; S2C-I**). As expected for this transgenic model, no significant changes in the expression of total APP were observed (**Figure 2G; S2C-D**). Semi-quantitative analysis of western blots showed a trending decrease in cortex expression of β-secretase (BACE-1) (**Figure S2F)** and anterior pharynx-defective phenotype 1 (APH-1) of the γ-secretase complex (**Figure S2I)** in 3xTg-AD x *Mcu*^fl/fl^ x Camk2a-Cre brains. We examined the expression of several APP processing fragments, including sAPPα, sAPPβ, AICD, and C99 and found no significant changes in expression (**Figure 2G; S2 J-M**). These results suggest that the reduction in Aβ levels observed in 3xTg-AD x *Mcu*^fl/fl^ x Camk2a-Cre mice is likely not due to alterative cleavage of APP. Instead, the decreased Aβ levels may result from changes in β-or γ-secretase activity or other indirect mechanisms affecting Aβ production or turnover. This implies that loss of MCU influences Aβ deposition through pathways other than direct alterations in APP processing.

**Figure 2.**
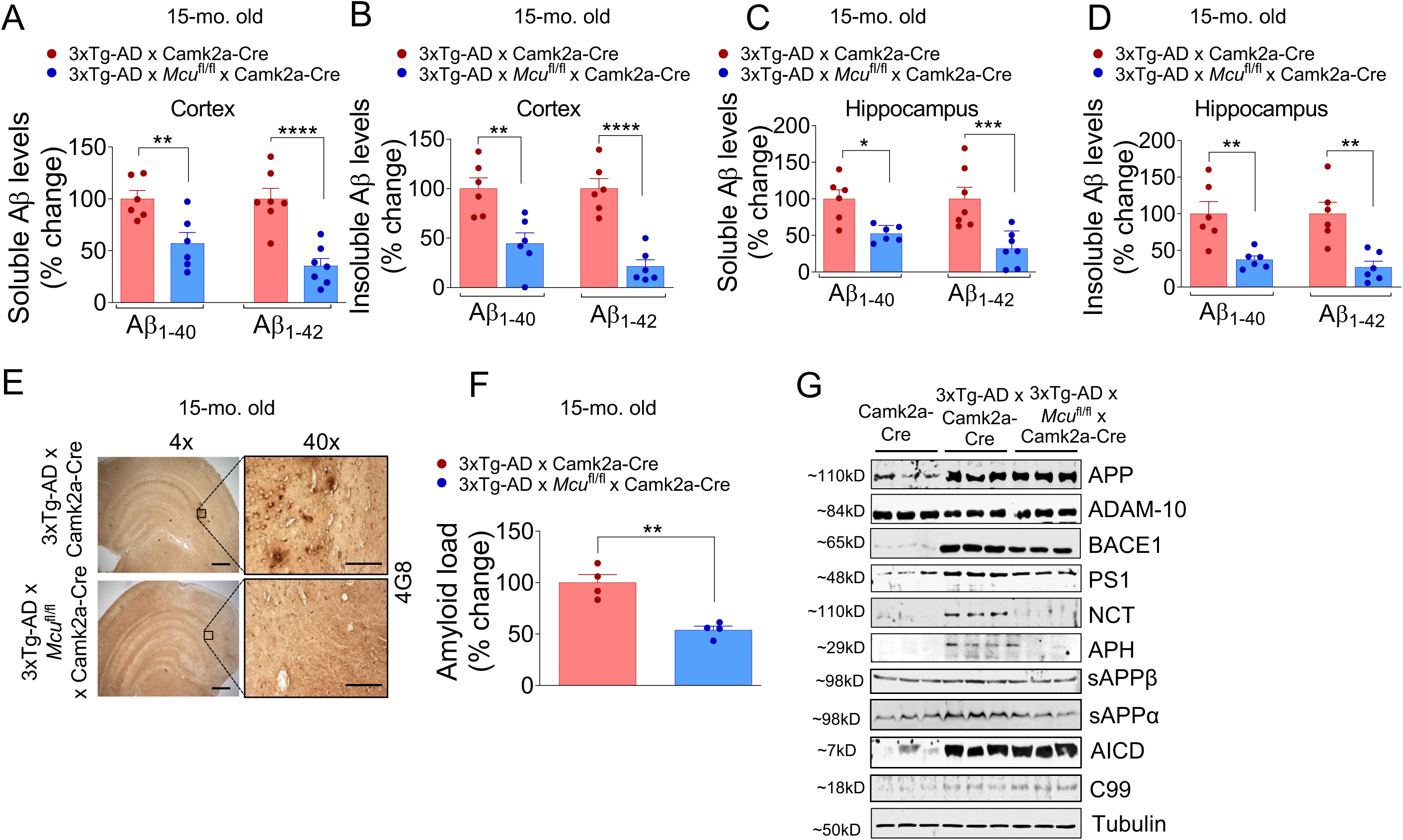
Genetic ablation of neuronal _m_Ca^2+^ uptake attenuates Aβ accumulation in 3xTg-AD mice. A, **B** Soluble and insoluble Aβ_1–40_ and Aβ_1–42_ levels in the cortex of 15 months old mice, measured by sandwich ELISA. **C, D** Soluble and insoluble Aβ_1–40_ and Aβ_1–42_ levels in the hippocampus of 15 months old mice, measured by sandwich ELISA. n = individual dots shown for each group in all graphs. **E** Representative immunohistochemical staining for 4G8-reactive β-amyloid; 4× scale bar = 100 μM, 40× scale bar = 50 μM. **F** Quantification of the integrated optical density area for Aβ immunoreactivity, n = 4 for all groups. **G** Western blots of full-length APP, ADAM-10, BACE1, PS1, Nicastrin, APH, sAPPα, sAPPβ, AICD, C99 and tubulin (loading control) for cortex homogenate of 15 months old mice, n = 3 for all groups. All data presented as mean ± SEM; ****p < 0.001, **p < 0.01; one-way ANOVA with Sidak’s multiple comparisons test.

To further investigate how MCU deletion impacts Aβ regulation, we examined the role of autophagy in clearing Aβ aggregates. We analyzed autophagy markers in protein lysates from the frontal cortex of 15-month-old control, 3xTg-AD, and 3xTg-AD x *Mcu*^fl/fl^ x Camk2a-Cre mice (**Figure S5 M-O**). However, we did not detect significant changes in the expression levels of LC3-II/LC3-I or p62, which are typically associated with autophagy activity. This could be due to the heterogenous tissue homogenate, which contains a significant number of non-neuronal cells that may mask differences in neuronal expression. Alternatively, these findings may suggest that the clearance of Aβ in this model does not occur through the traditional LC3-associated autophagy pathway and rather via alternative autophagic mechanisms.

### Loss of neuronal MCU expression decreases AD-associated tau pathology

Neurofibrillary tangles are a histopathological hallmark of AD and are predominantly comprised of hyper-phosphorylated tau. 15-month-old 3xTg-AD mice showed a significant increase in the expression of total soluble and insoluble tau, and increased phosphorylation at all residues examined compared to non-AD controls (**Figure 3A-F; S3A and B**). Deletion of *Mcu* from AD mice caused a significant reduction in soluble and insoluble tau (**Figure 3A-C; S3A**), a striking reduction in S202/T205 (AT8 immunoreactivity), T231/ S235 (AT180 immunoreactivity) and Ser396 (PHF-13 immunoreactivity) tau phosphorylation, (**Figure 3A, D-F; S3A)** and a variable reduction in T181 (AT270 immunoreactivity) (**Figure 3A;; S3A and B)**. To validate these findings we performed immunohistochemistry to examine tau phosphorylation specifically in pyramidal neurons of the CA1 region of the brain and found a significant reduction in S202/T205 (∼40%), T231/ S235 (∼34%), and PHF-13 (∼45%) tau phosphorylation in *Mcu*-deleted 3xTg-AD mice with no observed change in total soluble tau or AT270 (**Figure 3G-J; S3 C-D**). These results demonstrate that loss of neuronal _m_Ca^2+^ uptake is sufficient to reduce tau pathology in AD.

**Figure 3.**
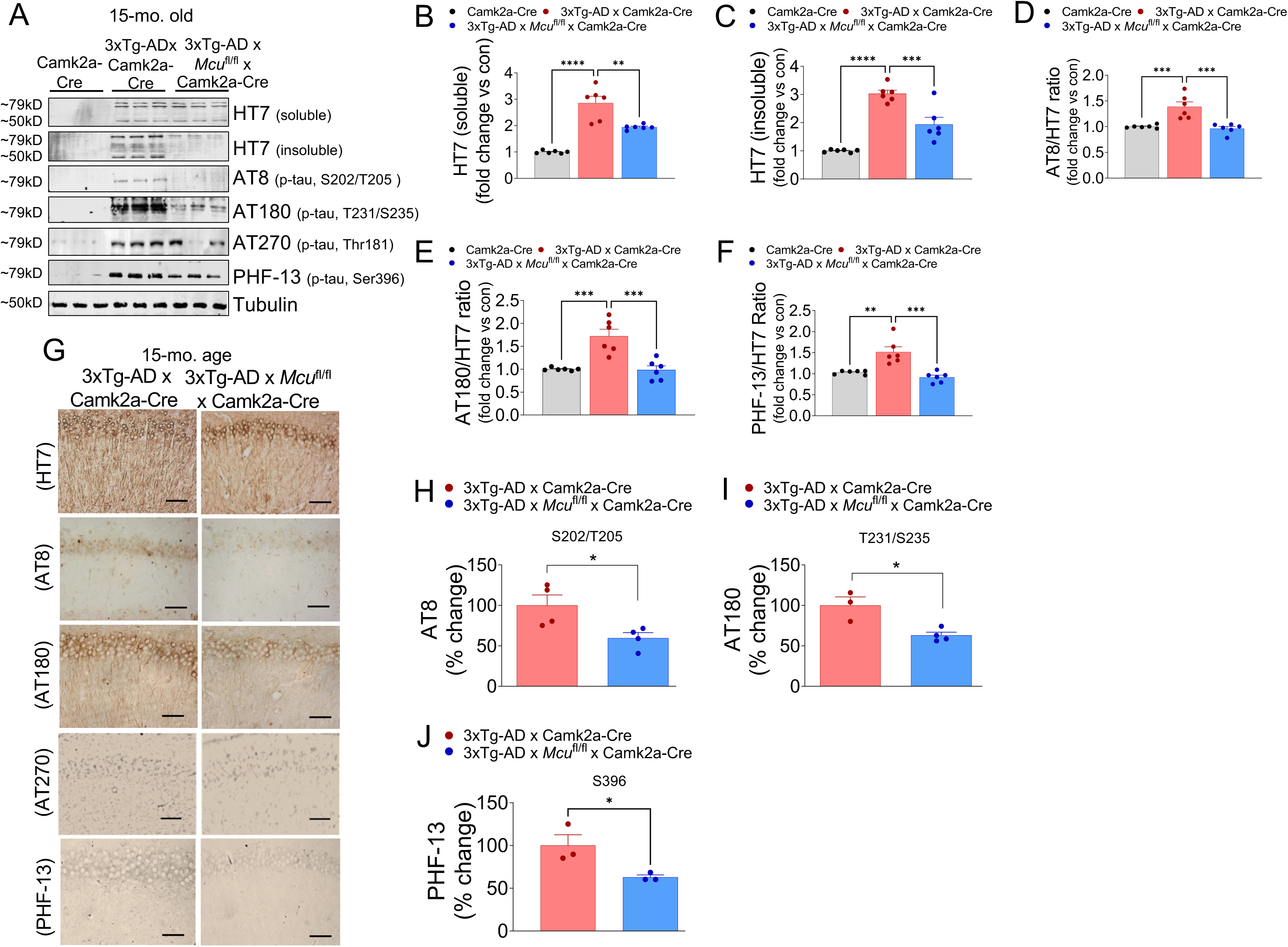
Loss of neuronal MCU expression decreases tau pathology. **A** Representative western blot of soluble and insoluble total tau (HT7), phosphorylated tau at residues S202/T205 (AT8), T231/S235 (AT180), T181 (AT270), and S396 (PHF13) in cortex homogenate of 15 months old mice, n = 3 for all groups. **B-F** Densitometric analysis of Western blots shown in Figure 3A and Figure S3A, expressed as fold-change vs. Camk2a-Cre con. corrected to a loading control tubulin. Quantification is the sum of all replicates, n=6. **G** Representative immunohistochemical staining for total tau (HT7), phospho-tau S202/T205 (AT8), phospho-tau T231/S235 (AT180) and S396 (PHF13) in the hippocampus of mice; scale bar = 50 μM. **H, I, J** Quantification of the integrated optical density area of AT8, AT180, AT270 and PHF-13 immunoreactivity, n = 4 for all groups.

To address the mechanism underlying reduced tau phosphorylation in MCU-deficient AD mice, we examined the levels of key tau kinases and phosphatases in protein lysate samples isolated from frontal cortex of 15-month-old Camk2a-Cre, 3xTg-AD x Camk2a-Cre, and 3xTg-AD x *Mcu*^fl/fl^ x Camk2a-Cre mice. We investigated the expression of GSK-3β, CDK5, ERK2, CaMKII, MARK, and RACK1 **(Figure S3 E-K)**. Increased GSK-3β expression has been associated with reduced neurogenesis and increased neuronal cell death in the hippocampus of AD patients (Llorens-Martin *et al*, 2013) . CDK5, a serine/threonine kinase, plays a role in neuronal survival and cognitive function by regulating synaptic morphology in the central nervous system. The CDK5/p25 complex, which is significantly elevated in AD patients, phosphorylates APP, Thr-252 of β-secretase, and STAT3, and is reported to exacerbate AD pathology (Cruz *et al*, 2003; Iijima *et al*, 2000). We found that the expression of GSK-3β and CDK5 was significantly increased in 3xTg-AD mice and expression was reduced in 3xTg-AD x *Mcu*^fl/fl^ x Camk2a-Cre mice (**Figure S3 E-G**). No significant changes were observed in the expression of MAPKs, CaMKII, MARK1, and ERK2 (**Figure S3 E; S3 H-K**).

We also examined the expression of major tau-related protein phosphatases, including Protein Phosphatase 2A (PP2A), Protein Phosphatase 1 (PP1), Protein Phosphatase 5 (PP5), and Calcineurin (PP2B), but found no significant changes (**Figure S3 L-P**) between 3xTg-AD x Camk2a-Cre and 3xTg-AD x *Mcu*^fl/fl^ x Camk2a-Cre mice. Overall, our comprehensive analysis highlights that while tau-related kinases show differential regulation, tau-related phosphatases remain unchanged. This sheds light on the molecular mechanisms by which preserved mitochondrial function may impact tau pathology and underscores its potential as a therapeutic target.

### Loss of neuronal mitochondrial calcium uptake reduces oxidative stress, improves synaptic integrity, and reduces neuroinflammation

Increased oxidative stress, neuroinflammation, and synaptic dysfunction are early cellular occurrences in AD pathogenesis (Butterfield & Boyd-Kimball, 2018; Hong *et al*, 2016). To examine the effect of neuronal loss of _m_Ca^2+^ uptake on redox status, we utilized dihydroethidium (DHE) to monitor real-time superoxide generation in fresh brain slices. Deletion of *Mcu* from 3xTg-AD mice caused a signficant reduction in ROS generation in both the cortex and hippocampus (**Figure 4A-C**). Next, we assessed lipid peroxidation by staining for 4-hydroxy-2-nonenal (4-HNE) in aged AD mice and controls. Deletion of *Mcu* from 3xTg-AD mice caused a ∼30% decrease in 4-HNE staining in the cortex and hippocampus (**Figure 4D-F**).

**Figure 4.**
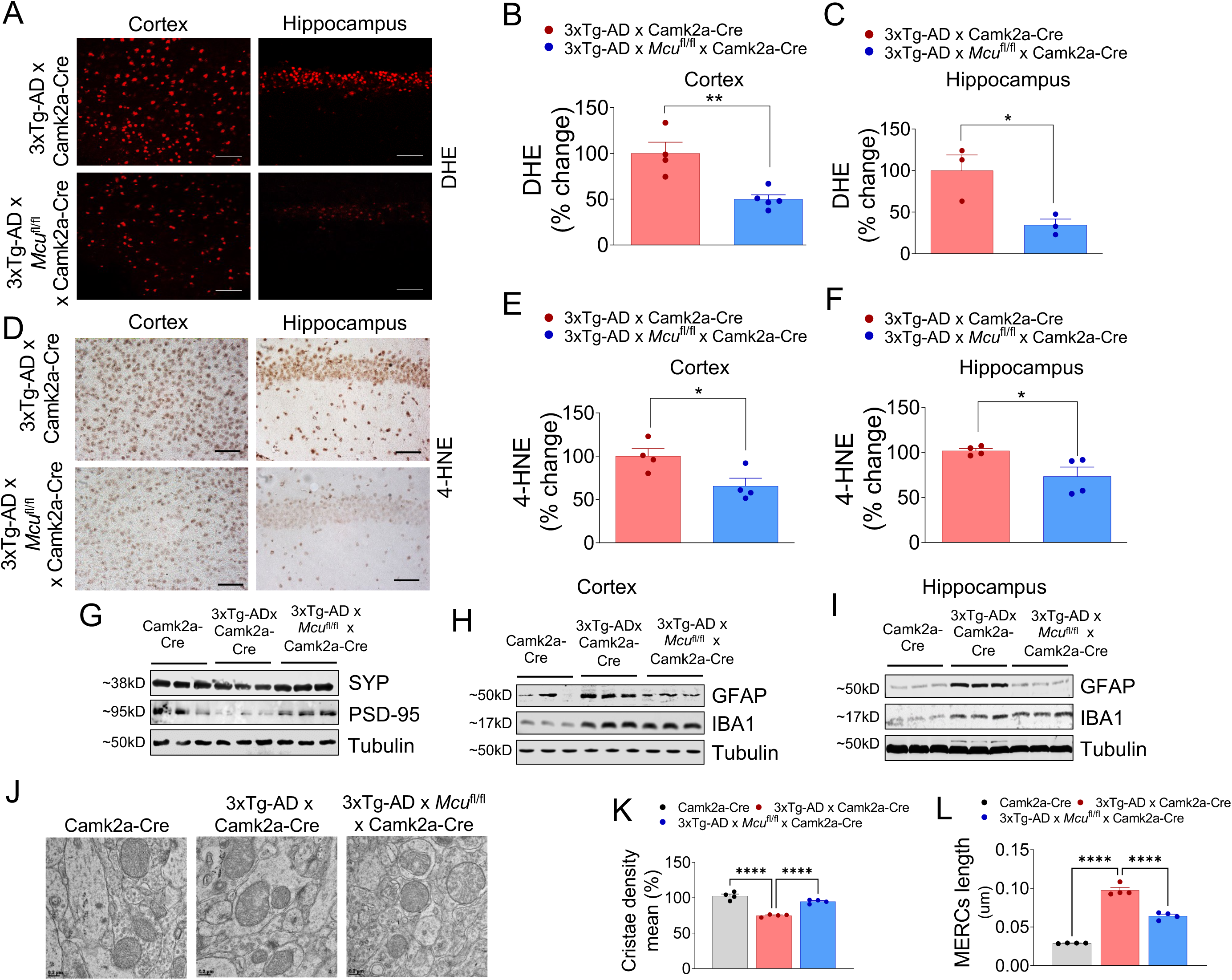
Neuronal deletion of *Mcu* reduces oxidative stress, improves synaptic integrity, and reduces neuroinflammation by preserving mitochondrial function. A DHE staining for ex vivo detection of superoxide production in freshly prepared cortical and hippocampal sections from 15 months old mice. **B, C** DHE fluorescent intensity, percent change vs. 3xTg-AD × Camk2a-Cre controls. n = 4 for all groups. **D** Representative image of 4-HNE immunohistochemistry in cortex and hippocampus to detect lipid peroxidation in 15 months old mice. **E, F** Percent change in 4-HNE-integrated optical density area corrected to 3xTg-AD × Camk2a-Cre controls. n = 4 for all groups. **G, H, I** Western blots for SYP, PSD-95, GFAP and IBA1 expression in tissue isolated from the cortex and hippocampus of 15 months old mice, n = 3 for all groups. **J** Representative transmission electron microscopy (TEM) images in the frontal cortex of 12-month-old mice; scale bar = 0.2 μM, n = 4 for all groups. **K** Cristae density (%) **L** Mitochondria-Endoplasmic reticulum contacts (MERC) distance (µM). All data presented as mean ± SEM; ****p < 0.001, ***p < 0.001, **p < 0.01, *p < 0.05; one-way ANOVA with Sidak’s multiple comparisons test.

Decreased synaptic plasticity and altered dendritic spines in excitatory glutamatergic synapses are early indicators of AD pathology (Almeida *et al*, 2005; Benarroch, 2018; Jackson *et al*, 2019). The vesicular protein, synaptophysin (SYP) is a marker of pre-synaptic integrity (Tarsa & Goda, 2002). Post-synaptic density protein (PSD-95) is a scaffolding protein localized at excitatory synapses (El-Husseini *et al*, 2000) which plays an important role in the process of learning (Migaud *et al*, 1998) and in the coupling of NMDA receptors and K^+^ channels at the post-synaptic membrane (Kim *et al*, 1996). Thus, we used these proteins as markers to assess synaptic plasticity and dendritic spine integrity. We observed a significant reduction in both SYP and PSD-95 expression in the brains of AD mice which was corrected to non-disease, control levels in the brains of 3xTg-AD x *Mcu*^fl/fl^ x Camk2a-Cre mice (**Figure 4G; S4A-B**). Similar findings were reported in (Cai H, 2022) where MCU knockdown increased the numbers of synapses and dendritic spines in 3xTg-AD mouse model.

Astrocyte and microglia activation are early features of AD-associated inflammation (Habib *et al*, 2020; Park *et al*, 2021). The expression of glial fibrillary acidic protein (GFAP), a marker of astrocyte reactivity (Caruso *et al*, 2013), was significantly decreased in brain cortex homogenates isolated from 3xTg-AD x *Mcu*^fl/fl^ x Camk2a-Cre, when compared to 3xTg-AD mice, without any changes in Ionized calcium binding adaptor molecule 1 (IBA1) expression, a marker of microglia activation (**Figure 4H; S4C-D**). Similarly, GFAP expression was significantly decreased, while IBA1 expression was unchanged, in the brain hippocampus of 3xTg-AD x *Mcu*^fl/fl^ x Camk2a-Cre, as compared to 3xTg-AD mice (**Figure 4I; S4E-F**). This suggests that the astrocyte mediated inflammatory response is rescued; however, there is no effect on microglia mediated inflammation in the hippocampus region of the AD mice brain with MCU inhibition.

To determine if changes in mitochondrial content occurred in our model system, we performed transmission electron microscopy (TEM) to examine *in situ* mitochondrial morphology. Four mice per genotype were examined and >200 mitochondria were analyzed per sample by a blinded investigator. No significant differences were observed in mitochondrial number per area or in mitochondrial shape (area, perimeter, circularity, Feret’s diameter, aspect ratio) in the frontal cortex of 12-month-old mice **(Figure 4J, S4 G-K)**. However, we observed significant changes in cristae density and mitochondrial apposition, the length of mitochondria and endoplasmic reticulum contacts (MERCs). Specifically, cristae density was increased in 3xTg-AD x *Mcu*^fl/fl^ x Camk2a-Cre mice compared to 3xTg-AD mice. Additionally, the mean MERC length was reduced in 3xTg-AD x *Mcu*^fl/fl^ x Camk2a-Cre mice, as compared to 3xTg-AD mice (**Figure 4 J-L**). The quantification of the ratio of mtDNA-to-nDNA corroborated that there was no change in mitochondrial content. In addition, there was no change in mitochondria mass between the groups at two timepoints (**Figure S4L**), but our data does support that mitochondrial mass decreases with age. All together, these results suggest that decreasing _m_Ca^2+^ uptake during AD pathogenesis reduces oxidative stress and inflammation and preserves synaptic integrity with improvements in mitochondrial morphology and sub-cellular apposition.

### Loss of _m_Ca^2+^ uptake prevents AD-associated mitochondrial dysfunction and promotes autophagic clearance of amyloid

To directly examine the mitochondrial consequences of *Mcu* loss and the molecular mechanisms affording neuroprotection, we utilized a neuroblastoma cell line harboring a transgene for the well characterized APP Swedish mutation (N2a-APPswe: K670N, M671L) (Thinakaran *et al*, 1996). We transduced control (N2a) and APPswe cells with lentivirus encoding shRNA targeting *Mcu* (**Figure 5A**). Western blot analysis revealed ∼55% loss of MCU protein in shRNA stable knockdown cells, as compared to scramble shRNA control cells (Scr-shRNA) (**Figure 5B, S5A**). We did not see any significant changes in the expression of other mtCU components in *Mcu* knockdown cells (**Figure S5B-D**). However, similar to what we previously reported in sporadic AD patients (Jadiya *et al*., 2019), we noted significant decreases in the expression of the mtCU regulators MICU1, MICU2, and MCUB (Mallilankaraman *et al*, 2012; Perocchi *et al*, 2010; Plovanich *et al*, 2013; Raffaello *et al*, 2013) in AD neurons and the loss of these proteins suggest a predisposition towards increased _m_Ca^2+^ uptake and overload (**Figure S5 B-D**). Attenuation of _m_Ca^2+^ uptake in *Mcu* knockdown cells was validated in a permeabilized cell system using the ratiometric reporters FuraFF (Ca^2+^) and JC1 (Δψ) (**Figure 5 C-D, S5F**). To evaluate matrix free-Ca^2+^ content, cells from all groups were permeabilized with digitonin and treated with thapsigargin to inhibit SERCA, and then treated with FCCP to release all matrix free-Ca^2+^, as previously reported (Jadiya *et al*., 2019; Luongo *et al*., 2015). APPswe cells were found to have increased _m_Ca^2+^ as compared to N2a controls. Further, knockdown of *Mcu* from APPswe cells significantly reduced _m_Ca^2+^ content (**Figure 5 E-F**).

**Figure 5.**
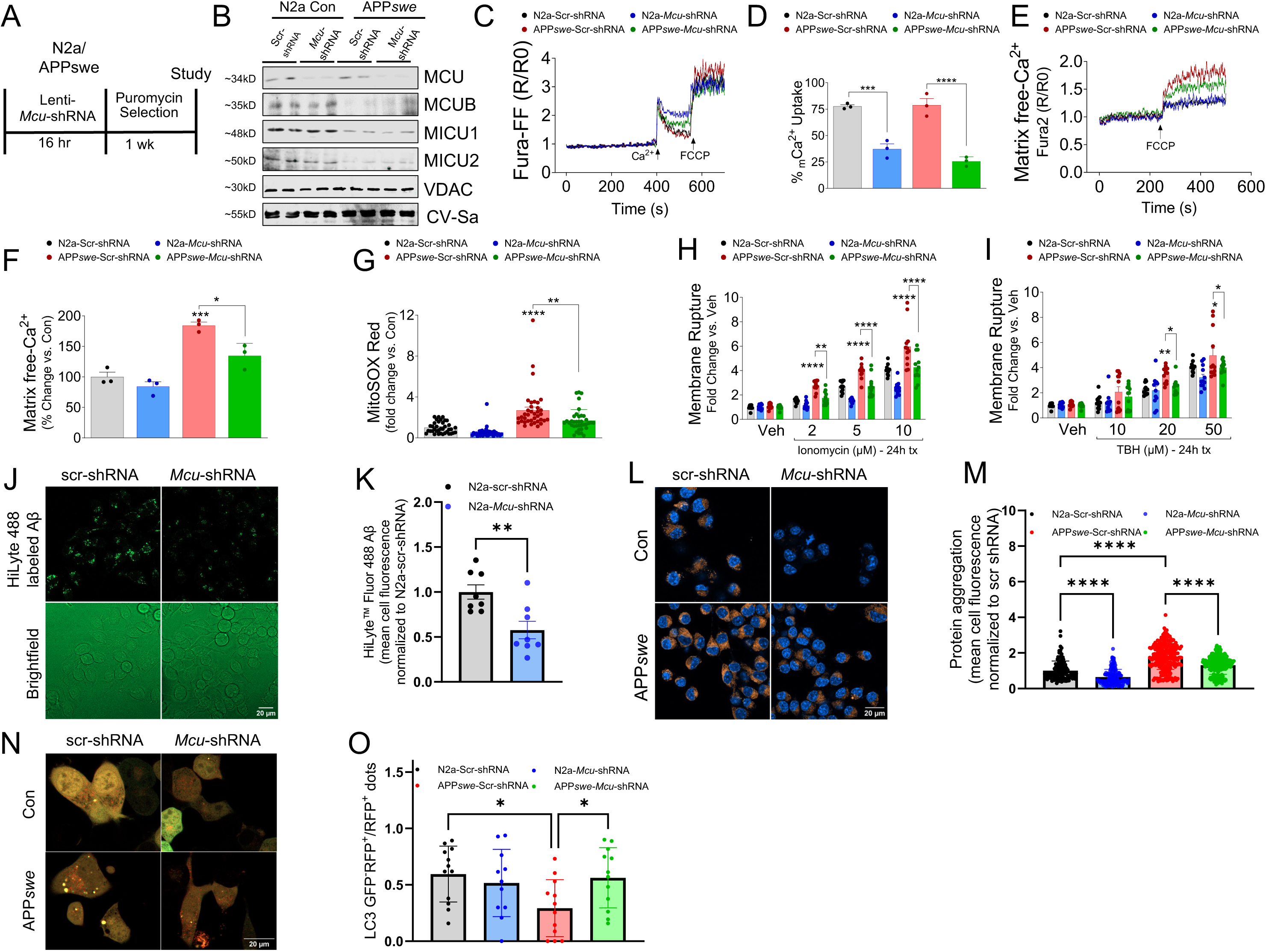
MCU loss in neurons prevents protein aggregation and amyloidogenic pathology by reducing mitochondrial ROS production and upregulating autophagy. **A** An Experimental protocol timeline for generation of stable Mcu knockdown N2a/APPswe cells. **B** Western blots for MCU expression and proteins associated with _m_Ca^2+^ uptake in N2a and APPswe cells transduced with lentivirus encoding shRNA targeting *Mcu.* C Representative recordings of _m_Ca^2+^ uptake. **D** Percent change in _m_Ca^2+^ uptake. controls **E** Representative traces for basal _m_Ca^2+^ content, n = 3. **F** Quantification of _m_Ca^2+^ content. **G** Quantification of MitoSOX fluorescent intensity; fold change vs. N2a Scr-shRNA controls. **H-I** Assessment for plasma membrane rupture, using Sytox Green after treatment with **H** Ionomycin (Ca^2+^ ionophore, 2–10 µM) and **I** tert-Butyl hydroperoxide (TBH, oxidizing agent, 10–50 µM). **J** Representative images for HiLyte™ Fluor 488-labeled Aβ 24 h after addition to the cells. **K** Quantification for J, normalized to scr-shRNA control. **L** Representative images of the fluorescent reporter Proteostat -protein aggregation. **M** Quantification for L, normalized to scr-shRNA control. **N** Representative images for cells expressing tandem fluorescent-tagged LC3 (mRFP-EGFP-LC3). **O** Quantification of autolysosome percentage to the total number of LC3 positive structures measured as ratio of LC3-GFP^-^RFP^+^ dots to LC3-RFP^+^ dots. n = individual dots shown for each group in all graphs. Data in **A-H** presented as mean ± SEM; Data in **J-O** presented as mean ± SD. To compare data in **J, K** t-test was used, for all the other data two-way ANOVA with Sidak’s multiple comparisons test. ****p < 0.001, **p < 0.01, *p < 0.05.

A number of studies suggest a strong association between _m_Ca^2+^ overload, oxidative stress and metabolic dysfunction (Gandhi *et al*, 2009; Luongo *et al*, 2017; Luongo *et al*., 2015; Sarasija *et al*., 2018) and both are hypothesized to be causal contributors to AD progression. We monitored the cells for changes in oxidative phosphorylation (OxPhos) by measuring mitochondrial oxygen consumption rates (OCR) in a Seahorse assay (**Figure S5 T-W**). Stable expression of mutant APPswe elicited a significant decrease in basal respiration and ATP-linked respiration and these parameters were not affected with loss of MCU (**Figure S5 T-W**). This result might be explained by a smaller role for mitochondrial calcium uptake in mitochondrial bioenergetics in immature neurons that mostly rely on glycolysis rather than oxidative phosphorylation. In support of this, APPswe cells displayed a more pronounced change in glycolysis, measured as extracellular acidification rate (ECAR), as opposed to OCR (**Figure S5U**). To examine redox status, we measured total cellular ROS content using the CellRox assay and mitochondrial superoxide production using the MitoSOX Red reporter. AD-mutant cell lines displayed enhanced ROS levels (total and mitochondrial), and both were reduced with loss of MCU (**Figure 5G, S5G**). To further evaluate MCU-mediated mechanisms of cytoprotection, cells from all groups were examined for plasma membrane rupture and viability following treatment with ionomycin (Ca^2+^ stress), and *tert*-butyl hydroperoxide (ROS stress). We observed a significant decrease in membrane rupture, a characteristic of necrotic cell death, and tendency toward enhanced general cellular viability in *Mcu* knockdown APPswe cell lines (**Figure 5 H-I, S5 H-I**).

To further define the molecular mechanisms affording cytoprotection against amyloid aggregation and AD pathology we extensively examined the gamma-secretase components, Nicastrin (NCT), Presenelin-1, and APH in N2a APPswe cells (**Figure S5 P-S**), but found no significant changes in expression. However, MCU loss did reduce Aβ accumulation 24-h after the addition of HiLyte™ Fluor 488-labeled Aβeta to the cells (**Figure 5 J-K**) and protein aggregation as measured by the Proteostat reporter dye (**Figure 5 L-M**) indicating a possible upregulation of clearance. Indeed, we found that APPswe reduces basal autophagy in N2a neurons, and this is rescued by MCU ablation measured as the percentage of autolysosomes corrected to the total number of LC3 positive structures in the resting condition using tandem fluorescent-tagged LC3 (mRFP-EGFP-LC3) (**Figure 5 N-O, S5 J-L**). Autophagic capacity can be impaired by the accumulation of damaged mitochondria thus creating a potential bottleneck in the clearance of amyloid and damaged material (Call & Nichenko, 2020; Nichenko *et al*, 2020). Autophagic capacity has also been reported to be reduced by mitochondrial dysfunction and excessive ROS generation (Vicente Roca-Agujetas, 2019). Importantly, all of these are partially rescued with the loss of MCU, providing a direct link between the inhibition of _m_Ca^2+^ uptake in neurons and protection against AD pathology.

## Discussion

Our group has previously reported that loss of _m_Ca^2+^ efflux capacity contributes to the pathogenesis and progression of Alzheimer’s disease (AD) by promoting mitochondrial Ca^2+^ (_m_Ca^2+^) overload. Here, we report that neuronal loss of mitochondrial Ca^2+^ uptake prevents AD-pathology and age-dependent cognitive decline in a robust mouse model of familial AD. While previous studies provide supportive anecdotal evidence of mitochondrial dysfunction resulting from _m_Ca^2+^ overload (Begley *et al*., 1999; Ferreira *et al*., 2015; Paula-Lima *et al*., 2011), our results demonstrate a causal role for dysregulated _m_Ca^2+^ transport mechanisms as an early pathogenic mechanism underlying the development and progression of AD and potentially other neurodegenerative diseases including ALS and Parkinson’s disease. In support, glutamate excitotoxicity due to excessive _i_Ca^2+^ promotes neuronal dysfunction in many diseases (Fan & Raymond, 2007; Lau & Tymianski, 2010; Wu *et al*, 2004), and MCU overexpression worsens excitotoxic cell death (Qiu *et al*., 2013). Previous studies demonstrate that _m_Ca^2+^ overload is sufficient to cause dendritic degeneration in a model of late-onset familial Parkinson’s disease (PD) (Verma *et al*., 2017). Further, pharmacologic and genetic inactivation of MCU is neuroprotective in a *pink1^−/−^* zebrafish model of PD (Soman *et al*, 2017; Soman *et al*, 2019). Strong correlative evidence that mtCU dysregulation is sufficient to cause brain and skeletal muscle disorders is seen in patients with loss of function mutations in *MICU1 (Logan et al., 2014)*. Indirect evidence from reports of increased ER-mitochondria crosstalk (Area-Gomez *et al*., 2012) and Ca^2+^ transfer in AD, also may contribute to increased _m_Ca^2+^ content. Indeed, neuronal cells expressing apolipoprotein E4 (apoE4), a major genetic risk factor for AD, displayed increased levels of _m_Ca^2+^ and ER-mitochondria tethering which promoted mitochondrial impairments and neuronal dysfunction (Orr *et al*, 2019). In totality, these reports coupled with our current results suggest alterations in mitochondrial Ca^2+^ handling are upstream of classical histopathological markers of AD such as Amyloid and Tao aggregation and deposition.

The exact mechanism for the neuroprotective effect of abrogating neuronal mitochondrial Ca^2+^ overload is likely multifactorial and involves decreasing mitochondrial ROS, preserving energetics, inhibiting cell death signaling, and enhancing autophagic capacity, ultimately preserving mitochondrial function. The AD brain is extremely vulnerable to redox imbalance due to the high neuronal energy demand. The increased expression and activity of β and γ secretase, and increased phosphorylation of tau are associated with oxidative stress (Jo *et al*, 2010), mitochondrial dysfunction (Gabuzda *et al*, 1994; Rhein *et al*, 2009), and impaired cellular energetics (Gabuzda *et al*., 1994). The link between Ca^2+^ and oxidative stress is supported by several previous reports. Elevated _i_Ca^2+^ enhances oxidative stress in neurons and astrocytes (Petersen *et al*, 2000). Moreover, the loss of MICU1 and a resultant increase in _m_Ca^2+^ content elicits mitochondrial superoxide generation (Mallilankaraman *et al*., 2012) and is linked with excitotoxicity (Starkov *et al*, 2004). In contrast, the inhibition of _m_Ca^2+^ uptake reduces NMDAR-induced excitotoxicity and neuronal cell death (Stout *et al*, 1998). Previous experiments support that inhibition of mtCU-dependent _m_Ca^2+^ uptake by either Ru360 treatment or siRNA knockdown of *Mcu* decreases oxidative stress in microglia cells *in vitro* (Xie *et al*, 2017) and in primary cerebellar granule neurons (Liao *et al*, 2015). Consistent with these findings, increased MCU activity is sufficient to promote oxidative stress (De Stefani *et al*., 2011; Liao *et al*., 2015). Moreover, increased ER-mitochondrial Ca^2+^ transfer in *PSEN* mutants is linked to high ROS production (Sarasija *et al*., 2018). Thus, as reported here, increased _m_Ca^2+^ content in AD can drive excess ROS production, furthering neuronal compromise and disease progression.

Another contributing mechanism underlying to the neuroprotective consequence of *Mcu* deletion may be reduced gliosis. Recently, MCU overexpression in cortical neurons of mice showed increased immunohistochemical staining for GFAP, suggestive of increased gliosis (Granatiero *et al*., 2019). Our current results agree with this observation, as we found a reduction in neuroinflammation with the ablation of _m_Ca^2+^ uptake. Furthermore, a consequence of reduced gliosis in AD may be preservation of synaptic integrity, as studies correlate enhanced gliosis with loss of PSD-95 (Gylys *et al*, 2004). However, the specific mechanisms by which defective _m_Ca^2+^ exchange modulates the functional synaptic unit and gliosis in AD remain to be elucidated.

Precisely why _m_Ca^2+^ uptake increases during AD pathogenesis remains unclear. Undoubtedly, neurons require a high energy supply for numerous functions such as synaptic transmission, action potential firing, and synapse development (Oyarzabal & Marin-Valencia, 2019; Vergara *et al*, 2019) and are highly dependent on mitochondrial metabolism to produce ATP (∼93%) via OxPhos (Harris *et al*, 2012). We hypothesize that neurons initially elevate _m_Ca^2+^ content via proteomic remodeling of transport machinery (NCLX, mtCU) to increase ATP production by increasing activity of the TCA cycle rate-limiting enzymes (PDH, α-KGDH, and ICDH). Proteomic remodeling of mtCU and NCLX were observed in our *in vivo* and *in vitro* AD models (Jadiya *et al*., 2019), suggesting that initially it may be an adaptive response to augment cellular energetics during stress, which subsequently becomes maladaptive and results in _m_Ca^2+^ overload, oxidative stress, and the accumulation of dysfunctional mitochondria causing a bottleneck in the clearance of damaged cell material by autophagy (Call & Nichenko, 2020; Nichenko *et al*., 2020). Altogether, this exacerbates amyloidosis and ultimately induces cell death. It is also known that aged or damaged mitochondria have a lower calcium buffering capacity and are more susceptible to calcium overload (Panel *et al*, 2018) which further supports mtCU dysregulation in AD. We previously demonstrated that _m_Ca^2+^ homeostasis can be restored by increasing NCLX _m_Ca^2+^ efflux in AD (Jadiya *et al*., 2019), and here we provide strong evidence that reducing mtCU _m_Ca^2+^ uptake lessens AD progression.

Even so, what is most impactful regarding this study remains the validation of mitochondrial calcium overload as a viable therapeutic target for halting the development and progression of AD. Despite significant investment and repeated attempts to leverage protein aggregation as a therapeutic strategy for AD, clinical trials on such drugs have been met with limited success and these drugs have not been widely adopted in the clinic due to limited benefit and safety concerns. Given the growing population of AD patients world-wide, there is an urgent need for novel therapeutic targets and strategies. Here, we have provided our second means of genetic proof-of-concept that mitochondrial calcium overload is a viable, safe, and effective strategy for the treatment of AD. Further studies are needed to determine the most efficacious pharmacological strategy for targeting _m_Ca^2+^ dysregulation in AD. Even so, our studies provided strong evidence that mitochondrial calcium is a causal contributor to AD pathogenesis and, most importantly, a promising therapeutic target in AD.

## Methods

### Neuronal specific *Mcu* knockout 3xTg-AD mutant mouse

*Mcu* floxed mice were generated as reported previously by our lab (Luongo *et al*., 2015) by acquiring targeted ES cells made by recombinant insertion of a construct containing loxP sites flanking exons 5-6 of the *Mcu* gene (ch10: 58930544-58911529). Mutant ES cell lines were confirmed by PCR and injected into C57BL/6N blastocysts with subsequent transplantation into pseudo-pregnant females. Germline mutant mice were crossed with ROSA26-FLPe knock-in mice for removal of the FRT-flanked neomycin resistance cassette. Resultant homozygous *Mcu*^fl/fl^ mice were crossed with neuron specific-Cre transgenic mice, *Camk2a*, to generate neuron-specific *Mcu* knockouts. The Calcium/calmodulin-dependent protein kinase II alpha (*Camk2a*) promoter drives Cre recombinase expression in the forebrain, specifically in the cortex and hippocampus (Tsien *et al*., 1996). Resultant neuronal-specific loss-of-function models (*Mcu*^fl/fl^ x Camk2a-Cre) were crossed with 3xTg-AD mutant mouse (Oddo *et al*, 2003) to generate 3xTg-AD x *Mcu*^fl/fl^ x Camk2a-Cre mutant mice. 3xTg-AD mice harbors three mutations: human Psen1 mutation (PS1^M146V^ knock-in), human amyloid precursor protein Swedish mutation (APP^Swe^ KM670/671NL), and P301L mutation of human tau (tau^P301L^).

### Cell cultures and stable cell lines

Mouse neuro-2a cells (N2a) and N2a cells stably expressing human APP with the K670N/M671L Swedish mutation (APPswe) were propagated in Dulbecco’s modified Eagle’s medium containing 10% fetal bovine serum and 1% penicillin/streptomycin. Cells were maintained at 37°C in a humidified 5% CO2 incubator. APPswe cells were grown in the presence of 400 µg/mL G418 (Jadiya *et al*., 2019). Differentiation of N2a and APPswe was performed using standard protocols described earlier (Evangelopoulos *et al*, 2005; Jadiya *et al*., 2019) in media containing 50% Dulbecco’s modified Eagle’s medium (DMEM) and 50% OPTI-MEM with 1% penicillin/streptomycin for 72 hrs (Jadiya *et al*., 2019). Cells were seeded onto poly-D-lysine coated glass coverslips for all imaging studies.

To generate *Mcu* knockdown stable N2a/APP cells, we transduced cells with lentivirus encoding shRNA targeting *Mcu* for 16 hrs. in the presence of 4mg/ml polybrene. Stably transduced cells were selected with puromycin (2 mg/ml) 48 hrs. post transduction for one week and expanded. Knockdown of *Mcu* was evaluated by Western blot.

### Immunohistochemistry

Mouse brains were dissected longitudinally at the center, and half of the brain was frozen on dry ice for biochemical analysis. The other was used for immunohistochemistry as previously described (Jadiya *et al*., 2019). In brief, brains were immersion-fixed in 4% paraformaldehyde (PFA) for 24 hrs. and embedded in paraffin. Serially sectioned (6-μm thick) brains were then deparaffinized, hydrated, and blocked in 2% fetal bovine serum. Sections were incubated overnight at 4 °C with following primary antibodies, monoclonal anti-β-amyloid, 17-24 (Aβ-4G8) dilution 1:150, HT7 dilution 1:150, phospho-tau (pThr231) monoclonal AT180 dilution 1:50, phospho-Tau (Ser202, Thr205) monoclonal AT8 dilution 1:50, anti-4 hydroxynonenal antibody (4-HNE) dilution 1:20, and then incubated with secondary antibody. The Vector Elite ABC system, an avidin/biotin-based peroxidase method, was used with diaminobenzidine, as the chromogen, to visualize immunoreactivity.

### Biochemical and Western blot analysis

Protein samples from frontal cortex and hippocampus of mouse brains as well as from cell lysates were lysed using a 1x RIPA lysis buffer with SIGMAFAST™ Protease Inhibitor Cocktail and phosphatase inhibitor for the soluble fractions. After ultracentrifugation at 90,720×*g* for 45 min at 4°C, the supernatant was kept as the soluble fraction. The pellet was then sonicated in the presence of 70% formic acid and ultracentrifuged at 90,720×*g* for 45 min at 4°C. The resulting formic acid-extracted supernatant representing the insoluble fraction was neutralized with 6 N sodium hydroxide. Aβ_1–40_ and Aβ_1–42_ levels both in soluble and insoluble fractions were assayed using the method described in the manual of Human Beta-Amyloid (1-40) ELISA Kit and Human Beta-Amyloid (1-42) ELISA Kit, respectively. The monoclonal antibody BAN50, specifically detects the N-terminal of human Aβ_1–16_, was used to capture Aβ_1–40_ and Aβ_1–42_ in samples. Captured Aβ_1–40_ and Aβ_1–42_ was recognized by BA27 F(Aβ’)2-HRP antibody, and BC05 F(Aβ’)2-HRP, respectively. Both BA27 and BC05 are mAβ that specifically detect the C-terminal of Aβ_._ The TMB based color development was used to assess the HRP activity, and absorbance was then measured at 450 nm. Values were presented as a percentage of Aβ_1–40_ and Aβ_1–42_ secreted relative to control.

For Western blot analysis, the protein samples were quantified using Bio-Rad Protein Assay Dye Reagent. Equal amounts of proteins were resolved by electrophoresis on SDS-PAGE gels and transferred to a PVDF Immobilon-FL membrane (EMD Millipore, Catalog # IPFL00010). The PVDF membrane was incubated with blocking buffer (Rockland, Catalog # MB-070) at room temperature for one hr. The membrane was incubated with primary antibody at 4°C overnight. The following primary antibodies were used in the study: MCU, MCUB, MICU1, MICU2, VDAC, ETC respiratory chain complexes, anti-APP N-terminal raised against amino acids 66–81 for total APP 22C11, BACE1, ADAM-10, PS1 (Sigma, P7854, 1: 500), Nicastrin (NCT, 3632S, cell signaling #14997, 1: 1000), APH1 (Millipore AB9214, 1:500), total tau HT7, phospho-tau (pThr231, AT180), phospho-Tau (Ser202, Thr205, AT8), phospho-tau (pThr181, AT270), phospho-tau (pS396, PHF13), beta-Tubulin, SYP dilution, PSD-95 dilution, GFAP. After incubation with primary antibody, the membrane was washed three times with TBS-T (TBS containing 0.1% Tween 20) for 10 min each and then incubated with specific secondary antibody for 1hr at room temperature. Licor IR secondary antibodies were used at dilutions 1:10,000. All blots were imaged on a Licor Odyssey system. All full-length western blots are available in Supplementary Figure 6.

### Memory tests

Mice at 6, 9, 12, and 15 months of age were assessed for spatial working memory in the Y-maze and hippocampal-dependent associative learning memory in fear conditioning assay. In our study, we used the same cohort of mice for the behavioral experiments, testing them at multiple time points. This longitudinal approach allows us to track changes within the same subjects over time, providing valuable insights into the progression of behavioral changes. The use of the same mice across different time points impacts the statistical analysis as it introduces intra-subject correlations. To account for this, we employed repeated measures ANOVA which is specifically designed for analyzing data from repeated measurements on the same subjects.

### Y-maze

We assessed spatial working memory by measuring spontaneous alternation in a Y-maze. In this assay, test animals were placed in the center of the Y-shaped maze (San Diego Instruments, 32 cm (long) 610 cm (wide) with 26-cm walls) for 5 min and the total number of arms entered, as well as the sequence of entries, were recorded. A spontaneous alternation was defined when a mouse enters a different arm of the maze in each of three consecutive arm entries (i.e., 1, 2, 3, or 2, 3, 1, or 3,1,2). Spontaneous alternation % was then calculated with the following formula: total alternation number/total number of entries-2 × 100. Prior to initial use, the arms of the maze were clearly designated as ’1’, ’2’ & ’3’, and the maze was always wiped clean with 70% ethanol, followed by water between each animal.

### Fear conditioning

The fear-conditioning was performed in a fear-conditioning apparatus (StartFear, Panlab Harvard Apparatus, 25 cm height × 30 cm width × 25 cm depth). The chamber was consisted of black methacrylate walls, a transparent front door, a light, a speaker, and a removable grid stainless-steel rod floor (3.2 mm diameter, 4.7 mm apart) through which a foot shock was administered. Automated fear conditioning FREEZING software was used to record and analyze signals generated by the animal movement throughout the procedure. Before each test, the chamber was cleaned with 70% ethanol, followed by water. In this test, mice were trained and tested on two consecutive days. On day 1 (training phase), each mouse was placed in the chamber and underwent three cycles of 30 s of sound and 10 s of electric shock (1.5 mA) within a 6-minute time interval. On day 2, the mouse returned to the same chamber without receiving electric shock or hearing the sound (contextual recall), and freezing behavior was recorded for 5 min. Two hours later, the animal spent 6 min in the same chamber but in an altered chamber environment, for example, different flooring, walls, smells, and lighting, and heard the cued sound for 30 s (cued recall), but without a foot shock. Freezing % was equal to (freezing time/total time) × 100%. All procedures were coordinated via PACKWIN (Panlab, Harvard Apparatus, USA) on a computer connected to the device, and data analysis was performed using the same software.

### Assessment of reactive oxygen species production

We employed Dihydroethidium (DHE) staining for in vivo detection of superoxide levels, as described (Jadiya *et al*., 2019). In brief, we freshly prepared cortical, and hippocampal slices from mice brains and stained them with 20 μM DHE for 30 min at 37°C and imaged at 518/excitation and 606/ emission on a Carl Zeiss 710 confocal microscope. To examine the total cellular ROS, cells from all groups were loaded with 5 μM CellROX Green Reagent for 30 min at 37 °C. CellROX exhibits a strong fluorogenic signal upon oxidation. The fluorescence at 485/excitation and 520/ emission was measured using a Tecan Infinite M1000 Pro plate reader. Next, we measured mitochondrial superoxide production using MitoSOX staining. Cells were loaded with 10 μM MitoSOX Red for 45 min at 37 °C and imaged at 490/20 excitation and 585/40 emission on Carl Zeiss 510 confocal microscope. All images were quantified for fluorescent optical density using ImageJ.

### Membrane Rupture and cell viability assay

We used SYTOX Green nucleic acid stain to examine membrane rupture and Cell Titer Blue to evaluate cell viability. SYTOX Green is a membrane-impermeable fluorescent stain that becomes permeable with compromised plasma membranes and intercalates in DNA and increases fluorescence. The Cell Titer Blue assay uses the indicator dye resazurin to measure cells’ metabolic capacity that an indicator of cell viability. In a 96-well plate, equal numbers of cells from all groups were treated with Ionomycin (2–10 µM) and an oxidizing agent tert-Butyl hydroperoxide solution (TBH: 10–50 µM) for 24 hrs. On the day of the experiment, SYTOX green dye was added to wells at 1 μM final concentration for 15 min at 37 °C. Fluorescence was measured using a Tecan Infinite M1000 Pro plate reader at 504/ excitation and 523/ emission. To measure the number of viable cells, Cell Titer Blue Reagent (10µl/well in 96 well plates) was added directly to each well and incubated at 37°C for 2 hrs. Fluorescence was measured using a Tecan Infinite M1000 Pro plate reader at 560/ excitation and 590/ emission. Data were normalized to vehicle control to avoid any differences in cell numbers between the groups.

### Evaluation of _m_Ca^2+^ retention capacity and content

Assessment of _m_Ca^2+^ uptake and content were performed as reported previously (Jadiya *et al*., 2019). Cells (2 × 10^6^) from all groups were washed in Ca^2+^ -free DPBS and resuspended in an intracellular-like medium (120 mM KCl, 10 mM NaCl, 1 mM KH2PO4, 20 mM HEPES-Tris) containing thapsigargin (3 μM) to block the SERCA pump, digitonin (80-μg/ml), protease inhibitor, and succinate (10 μM) at pH 7.2. All solutions were cleared with Chelex 100 to remove trace Ca^2+^. To measure _m_Ca^2+^ uptake, cells were gently stirred and loaded with the ratiometric reporters Fura-FF at concentration of 1 μM to monitor extra-mitochondrial Ca^2+^. To monitor mitochondrial membrane potential (Δψ), JC1 was added at 20 s. Fluorescence signals were monitored at 340-and 380-nm excitation /510-nm emission for Fura-FF to calculate ratiometric changes and at 490 nm excitation/535 nm emission for the monomer 570 nm excitation/595 nm emission for the J-aggregate of JC-1. After acquiring baseline recordings, a 10 µM Ca^2+^ bolus was added at 400 s. Clearance of extra-mitochondrial Ca^2+^ was representative of _m_Ca^2+^ uptake. At 550s, a protonophore, 10 μM FCCP (a protonophore) was added to uncouple the Δψ_m_ and release matrix-free-Ca^2+^. All experiments (three replicates) were conducted at 37°C and recorded on a PTI spectrofluorometer (Delta RAM, Photon Technology Int.). The _m_Ca^2+^ uptake rate was recorded over 50s post Ca^2+^ bolus. To evaluate _m_Ca^2+^ content, permeabilized cells from all the groups were loaded with Fura-2 for ratiometric monitoring of extra-mitochondrial Ca^2+^ using a fluorescent spectrofluorometer. Upon reaching a steady-state recording, FCCP was used to collapse ΔΨ_m_ and liberate all matrix-free-Ca^2+^. To measure basal _m_Ca^2+^ in intact cells, cells were transfected with mitochondrial genetically encoded calcium indicator, CMV-mito-GEM-GECO1 (Addgene, plasmid #32461) using FuGENE HD (Promega), and imaged 48 h after using a Carl Zeiss 900 confocal microscope (405 nm laser excitation and blue (470 nm) and green (535 nm) emission). Obtained images were analyzed in Fiji 2.14.0 using custom macros (https://sites.imagej.net/Theweave) for background subtraction. Individual cells were masked and mean gray values were exported and further processed in Excel and GraphPad Prism 10.4.0. _m_Ca^2+^ was calculated as the ratio of blue to green emission.

To assess _m_Ca^2+^ retention capacity in vivo, we next isolated mitochondria from frontal cortex of mice and performed Calcium Green 5 n to monitor extra-mitochondrial Ca^2+^ as described previously (Jadiya *et al*., 2019). After baseline recordings at 400 s, 5 µM-repetitive Ca^2+^ boluses were added and fluorescent signal at (488Ex/530Em) was measured using plate reader.

### Autophagy assay

To estimate basal autophagy, tandem-tagged LC3-EGFP-RFP assay was used. Cells were transfected with tandem-tagged LC3-EGFP-RFP (Addgene, plasmid #21074) using FuGENE HD (Promega), and imaged 48 h after using a Carl Zeiss 900 confocal microscope using GFP and RFP settings. Obtained images were used to count the number of GFP^+^RFP^+^ and GFP^-^ RFP^+^ dots per cell representing autophagosomes and autolysosomes, respectively. Percentage of autolysosomes to the total number of dots were calculated.

### Estimation of Aβ accumulation

To estimate amyloid accumulation, HiLyte™ Fluor 488-labeled Aβ (Anaspec, AS-60479-01) was added to cell media at 1 µM. The cells were examined 24 h after using a Carl Zeiss 900 confocal microscope using GFP settings. Obtained images were analyzed in Fiji 2.14.0 using custom macros (https://sites.imagej.net/Theweave) for background subtraction. Individual cells were masked and mean gray values were exported and further processed in Excel and GraphPad Prism 10.4.0.

### Detection of protein aggregates

Protein aggregates were detected as described previously (Jadiya *et al*., 2019). Briefly, for determination of misfolded protein aggregates, cells were fixed with 4% paraformaldehyde at RT for 15 min and, permeabilized in PBST (0.15% TritonX-100 in PBS) at RT for 15 min. Cells were then stained with Proteostat aggresome detection dye at RT for 30 min and Hoechst 33342 nuclear stain, using the method described in the manual (Enzo Life Science Inc., Farmingdale, NY, USA). Proteostat (Enzo), a molecular rotor dye that becomes fluorescent when binding to the β-sheet structure of misfolded proteins. All components of the Proteostat aggresome detection kit were prepared according to the manufacturer’s instructions. Aggregated protein accumulation was detected using a Carl Zeiss 900 confocal microscope (standard red laser set for the aggresome signal and DAPI laser set for the nuclear signal imaging). Obtained images were analyzed in Fiji 2.14.0 using custom macros (https://sites.imagej.net/Theweave) for background subtraction. Individual cells were masked and mean gray values were exported and further processed in Excel and GraphPad Prism 10.4.0.

### Mitochondrial bioenergetics

Mitochondrial bioenergetics was estimated as described previously (Jadiya *et al*., 2019). Briefly cells were subjected to OCR measurement at 37 °C in an Agilent Seahorse XF Pro analyzer (Agilent). Cells (4 × 10^4^) were plated in regular medium and next day assayed in bicarbonate-free phenol red-free DMEM media (corning 90-113-PB) supplemented with 10 mM glucose, 2 mM L-glutamine, 1 mM sodium pyruvate, and 5 mM HEPES pH 7.4. During the assay cells were sequentially exposed to oligomycin (2 µM), FCCP (2 µM), and rotenone plus antimycin A (1 µM). Quantification of basal respiration (base OCR – non-mito respiration (post-Rot/AA), ATP-linked respiration (post-oligo OCR−base OCR), Max respiratory capacity (post-FCCP OCR−post-Rot/AA), Spare respiratory capacity (post-FCCP OCR−basal OCR) and Proton leak (post-Oligo OCR−post-Rot/AA OCR) was performed.

### Transmission electron microscopy

Mice were anesthetized and perfused with PBS for 5 min and then with a fixative solution of 2% paraformaldehyde/2% glutaraldehyde in 0.1M sodium cacodylate buffer. After dissection, intact brain was placed in a fresh solution of the fixative overnight at 4 °C under agitation. Coronal sections (100-200um) were cut with the vibratome and then processed the tissue slices and embed in epoxy resin over a period of 48 h. The areas of interest (cortex, three blocks per mouse) were cut out of the slices with Reichert-Jung Ultracut E ultramicrotome and mounted for ultrathin sectioning for the TEM on Athene 200 mesh thin bar formvar-carbon coated copper grids. Sections were visualized with a Zeiss Libra 120 transmission electron microscope operating at 120 kV, using a Gatan Ultrascan 1000 CCD camera. We used ImageJ software to analyze mitochondria ultrastructure and mitochondrial shape descriptors (area, perimeter, circularity, Feret’s diameter, aspect ratio). Minimum of 200 mitochondria were analyzed per mouse, *n* = 4 mice per group.

### Quantification of mitochondrial copy number

The genomic DNA was isolated from mice brain cortex tissue using the DNeasy blood and tissue kit (Qiagen 69504) according to manufacturer’s instructions. Quantification of mtDNA copy number was performed via quantitative PCR (qPCR) using the primers mitochondrial COXII gene and nuclear β-Globin (Jadiya *et al*., 2019).

### Statistics

Prism 6.0 Graph Pad software was used for statistical analysis and to generate plots. All results are shown as mean ± SEM. All experiments were repeated at least three times and measurements were taken on distinct samples. Individual data points mean and s.e.m. were displayed in the figures. Where appropriate column analyses were performed using an unpaired, two-tailed *t* test (for two groups) or one-way ANOVA with Bonferroni correction (for groups of three or more). For grouped analyses either multiple unpaired t-test with correction for multiple comparisons using the Holm–Sidak method or where appropriate two-way ANOVA with Tukey post hoc analysis was performed. The specific statistical tests for each experiment are indicated throughout the manuscript. Results were significant if p values < 0.05 (95% confidence interval).

### Study approval

Animal studies were approved by Temple University’s IACUC and followed AAALAC guidelines.

## Author contributions

J.W.E. conceived the project; J.W.E., P.J., and E.B. contributed to study design, data analysis and writing the paper; P.J., E.B., D.T., D.K., S.S., M.T., L.M., A.N.H. and S.K. performed experiments, data collection and interpretation, P.J. and D.K. performed histology and memory test experiments, D.T. and M.T. assisted with calcium content and imaging assays; J.W.E., J.G., E.B., and H.C. edited the manuscript and provided expertise with data interpretation.

## Acknowledgments

We thank the late Trevor Tierney for technical assistance in the Elrod Lab. Support for this work was provided by grants from the NIH: R01NS121379, R01HL136954, R01HL142271, 3R01HL123966-05S1, P01HL147841, P01HL134608; AHA: 20EIA35320226, and Pennsylvania Dept. of Health CURE Award (420792) to J.W.E.; NIH K99AG065445, Alzheimer’s Association 24AARG-D-1191292, American Heart Association 24IPA1273195, Wake ADRC REC and Development grant (P30AG072947) to P.J., NIH K99DK120876 to D.T. and NIH F32HL151146 to J.F.G.

## Supplementary material

Supplementary information including supplementary figures and can be found online with this article.

## Supplementary Information

Genetic ablation of neuronal mitochondrial calcium uptake impedes Alzheimer’s disease progression Jadiya et al.

---

**Supplementary Figure 1.**
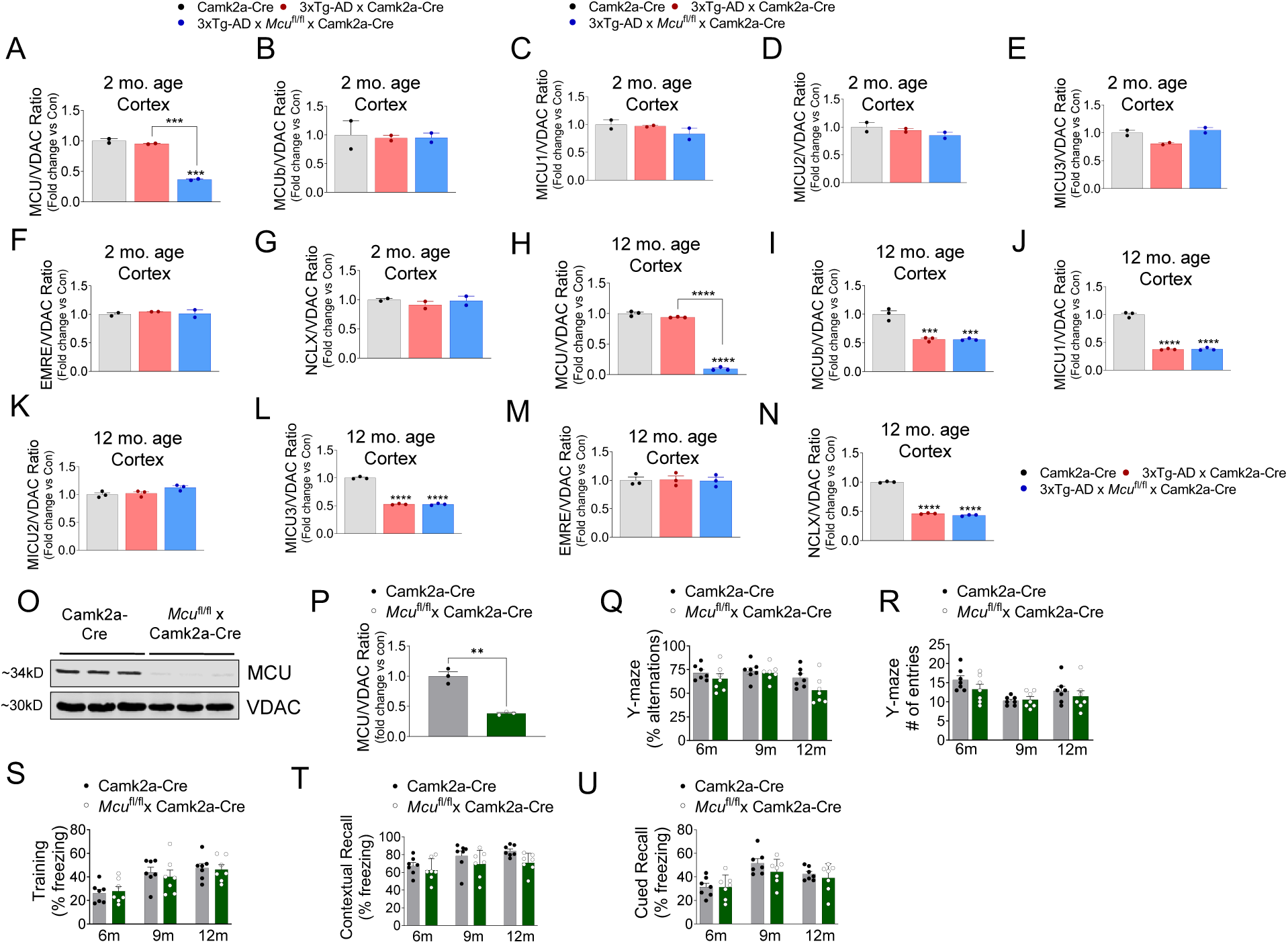
_m_Ca^2+^ exchanger expression and cognitive assay. **A-N** Quantification of protein expression associated with _m_Ca^2+^ exchange expressed as fold-change vs. Camk2a-Cre con. corrected to a mitochondrial loading control VDAC, in tissue isolated from the brain cortex of 2-and 12-month-old mice. **O, P** Western blot validation and densitometry analysis for the expression of MCU protein in tissue isolated from the cortex of *Mcu*^fl/fl^ x Camk2a-Cre mice compared to age-matched control corrected to VDAC**. Q, R** Y-maze spontaneous alternation test. **Q** Percentage of spontaneous alternation. **R** Total number of arm entries. **S-U** Fear-conditioning test. **S** Freezing responses in the training phase. **T** Contextual recall freezing responses, **U** Cued recall freezing responses. n = individual dots shown for each group in all graphs. All data presented as mean ± SEM; ****p < 0.001, ***p < 0.001, *p < 0.05; one-way ANOVA with Sidak’s multiple comparisons test.

**Supplementary Figure 2.**
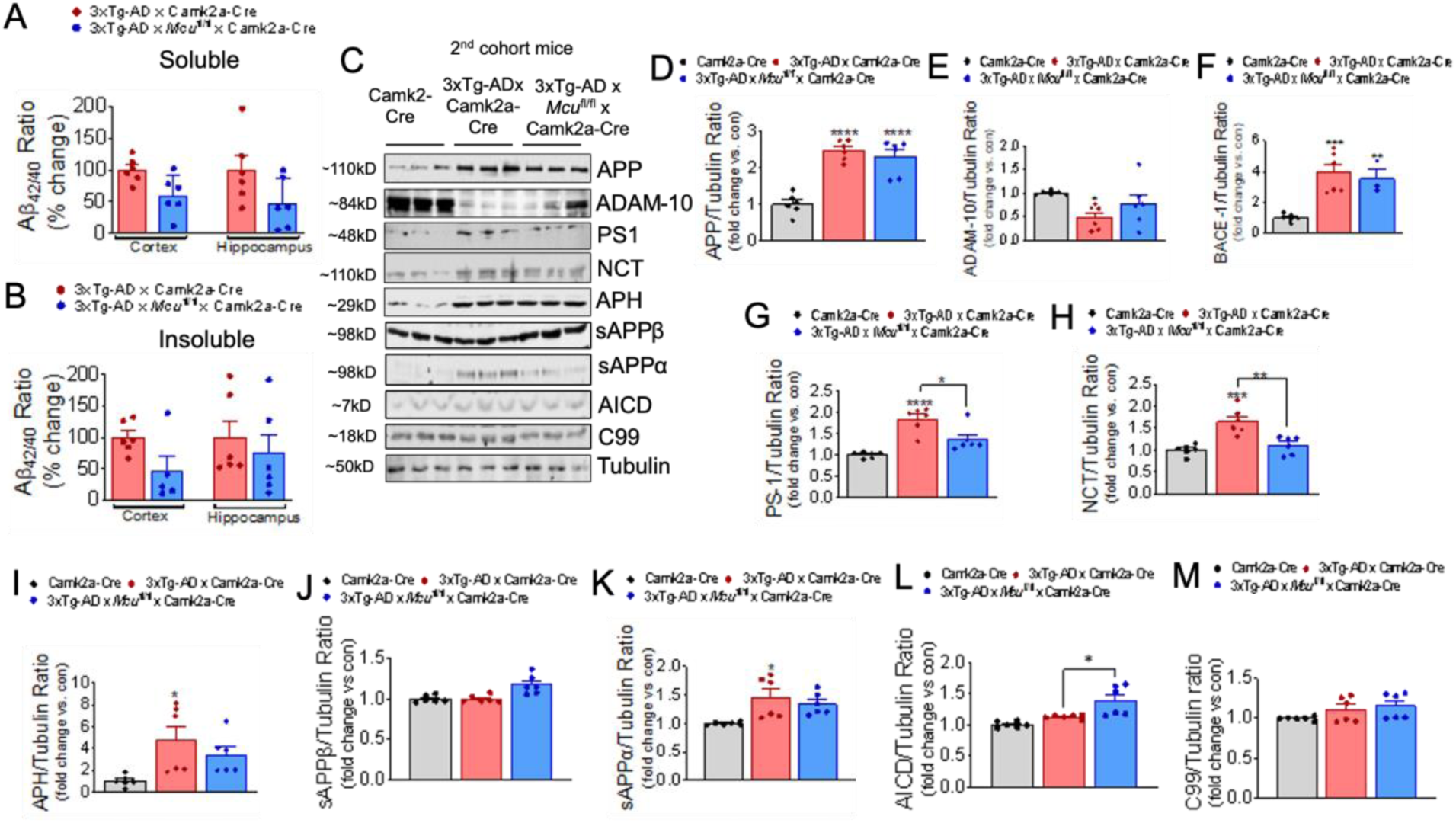
Effect of genetic ablation of neuronal _m_Ca^2+^ uptake on Aβ pathway. **A** Soluble Aβ_1–42_/Aβ_1–40_ ratio in cortex and hippocampus of 15-month-old mice, measured by sandwich ELISA. **B** Insoluble Aβ_1–42_/Aβ_1–40_ ratio in cortex and hippocampus of 15-month-old mice, measured by sandwich ELISA. **C** Western blots of full-length APP, ADAM-10, PS1, nicastrin, APH, sAPPα, sAPPβ, AICD, C99 and tubulin (loading control) for cortex homogenate of 15-month-old mice (second cohort mice), n = 3 for all groups. **D-M** Densitometry analysis of Western blots shown in Figure 2G and Figure S2C, expressed as fold-change vs. Camk2a-Cre con. corrected to a loading control tubulin. n = individual dots shown for each group in all graphs. All data presented as mean ± SEM; ****p < 0.001, **p < 0.01, *p < 0.05; one-way ANOVA with Sidak’s multiple comparisons test.

**Supplementary Figure 3.**
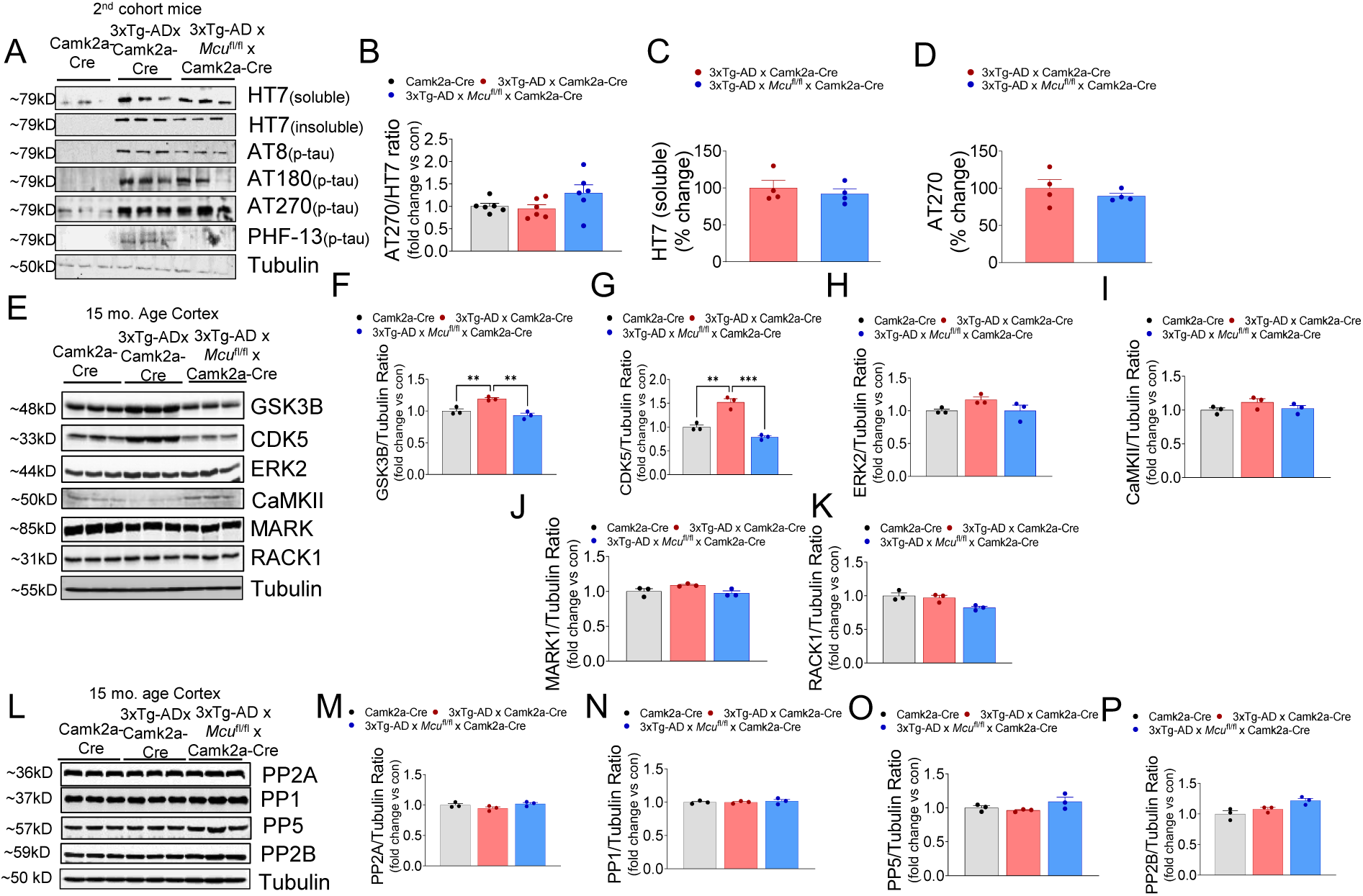
Effect of loss of neuronal MCU expression on tau pathology. **A** Western blots (2^nd^ cohort mice) of soluble and insoluble total tau (HT7), phosphorylated tau at residues S202/T205 (AT8), T231/S235 (AT180), T181 (AT270), and S396 (PHF13) in cortex homogenate of 15-month old mice, n = 3 for all groups. **B-D** Densitometric analysis of Western blots shown in Figure 3A, and Figure S3A, expressed as fold-change vs. Camk2a-Cre con. corrected to a loading control, tubulin. Quantification is the sum of all replicates, n=6. **E** Western blots for protein expression of GSK3β, CDK5, ERK2, CamkII, MARK, RACK1 and tubulin (loading control) for cortex homogenate of 15 months old mice, n = 3 for all groups **K-N** Densitometric analysis of western blots shown in Figure S3J, expressed as fold-change vs. Camk2a-Cre con. corrected to a loading control, tubulin. **L** Western blots for protein expression of PP2A, PP1, PP5, PP2B and tubulin (loading control) for cortex homogenate of 15 months old mice, n = 3 for all groups **M-P** Densitometric analysis of western blots shown in Figure S3L, expressed as fold-change vs. Camk2a-Cre con. corrected to a loading control, tubulin. All data presented as mean ± SEM; ****p < 0.001, **p < 0.01, *p < 0.05; one-way ANOVA with Sidak’s multiple comparisons test.

**Supplementary Figure 4.**
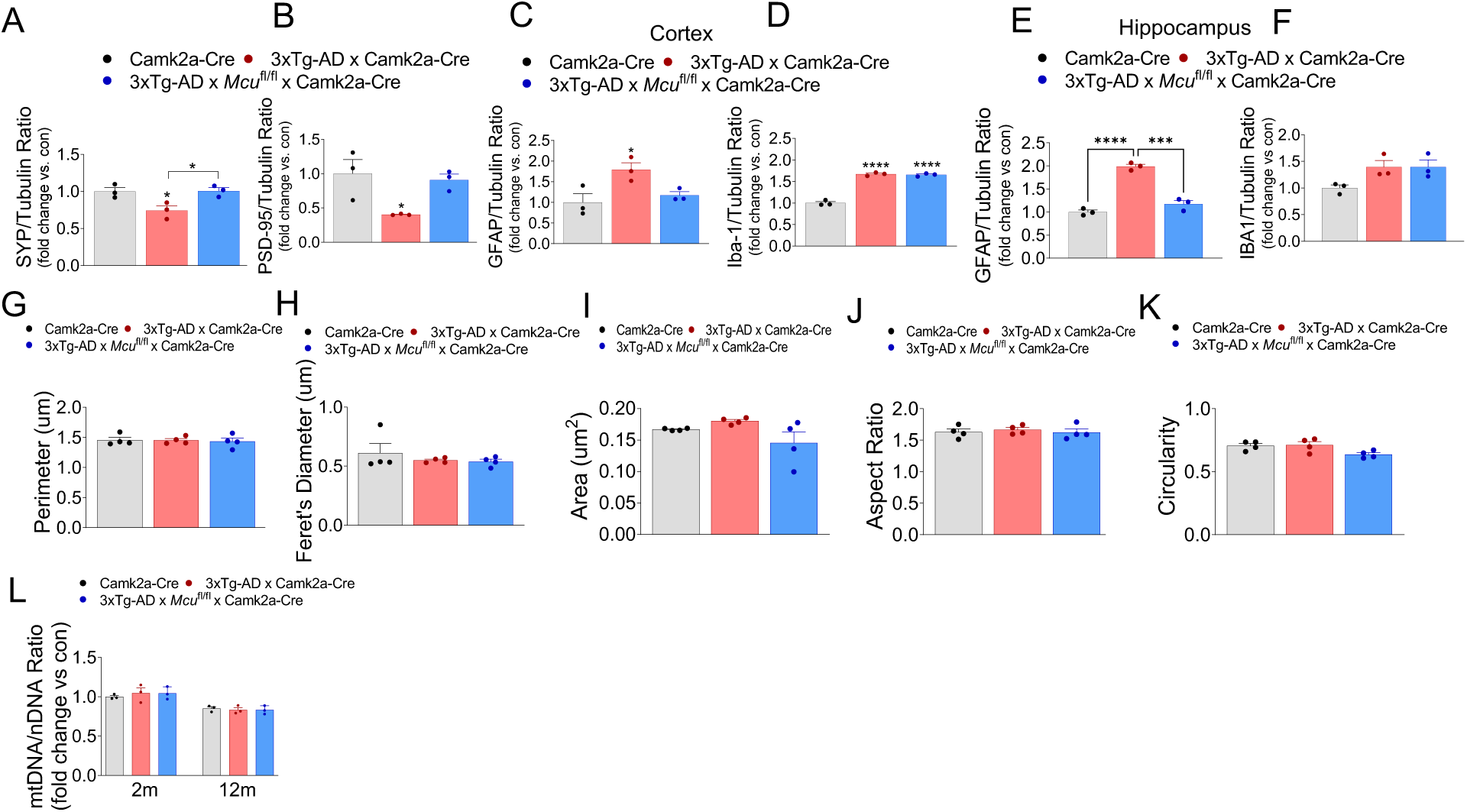
A-F Densitometric analysis of Western blots shown in Figure 4 G, H, I for SYP, PSD-95, GFAP and IBA1 expression. n = 3 for all groups. **G-K** Quantification of the shape descriptors and morphological parameters for mitochondria. **G** Perimeter (μM). **H** Feret’s diameter (μM). **I** Area (μM^2^). **J** Aspect ratio. **K** Circularity. **L** Mitochondrial DNA (mtDNA)/nuclear DNA (nDNA) ratio in tissue isolated from the cortex of 2-and 12-months old mice, fold change vs. 2 months old Camk2a-Cre controls. All data presented as mean ± SEM; ****p < 0.001, ***p < 0.001, **p < 0.01, *p < 0.05; one-way ANOVA with Sidak’s multiple comparisons test.

**Supplementary Figure 5.**
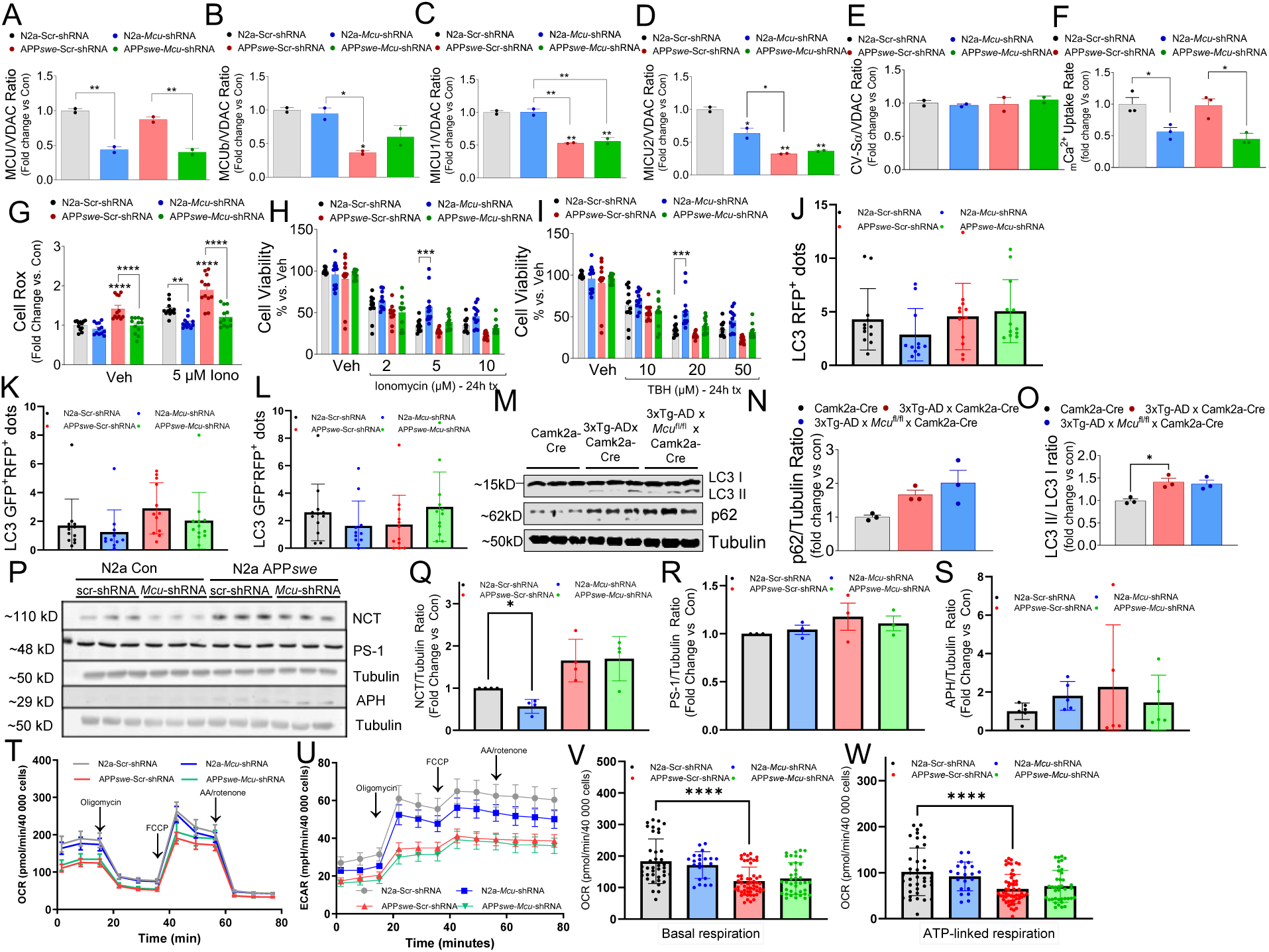
A-E Quantification of protein expression associated with _m_Ca^2+^ exchange expressed as fold-change vs. N2a Scr-shRNA con. and corrected to a mitochondrial loading control, VDAC. **F** Fold-change in _m_Ca^2+^ uptake rate of N2a-*Mcu*-shRNA, APPswe and APPswe -*Mcu*-shRNA vs. N2a Scr-shRNA controls. **G** Quantification of CellRox green fluorescent intensity (total cellular ROS production); fold change vs. N2a Scr-shRNA controls. **H, I** Assessment for cell viability using Cell Titer Blue after treatment with, **H** Ionomycin (Ca^2+^ overload, 2-10 µM), **I** tert-Butyl hydroperoxide (TBH, oxidizing agent, 10-50 µM), n = individual dots shown for each group. **J-L** Quantification of autophagic structures per cell, including: **J** RFP⁺ puncta representing total red puncta **K** GFP⁺RFP⁺ puncta representing autophagosomes **L** GFP⁻RFP⁺ puncta representing autolysosomes. **M** Representative western blot of LC3-I, LC3-II, p62 and tubulin (loading control) for cortex homogenate of 12-month-old mice, n = 3 for all groups. **N, O** Densitometry analysis of Western blots shown in Figure S2M expressed as fold-change vs. Camk2a-Cre con. corrected to tubulin n = 3/groups. **P** Western blots for protein expression of NCT, PS-1, APH and tubulin (loading control) for cells homogenate of N2a Scr-shRNA, N2a-*Mcu*-shRNA, APPswe and APPswe -*Mcu*-shRNA. **Q-S** Densitometric analysis of western blots shown in Figure S5P, expressed as fold-change vs. N2a Scr-shRNA con. corrected to a loading control, tubulin. **T–W.** Metabolic profiling of neuronal cells. **T** Representative traces of oxygen consumption rate (OCR) analysis **U** Representative extracellular acidification rate (ECAR) traces **V–W,** Quantification of basal respiration and ATP-linked respiration in N2a Scr-shRNA, N2a Mcu-shRNA, APPswe, and APPswe Mcu-shRNA

**Supplementary Figure 6.**
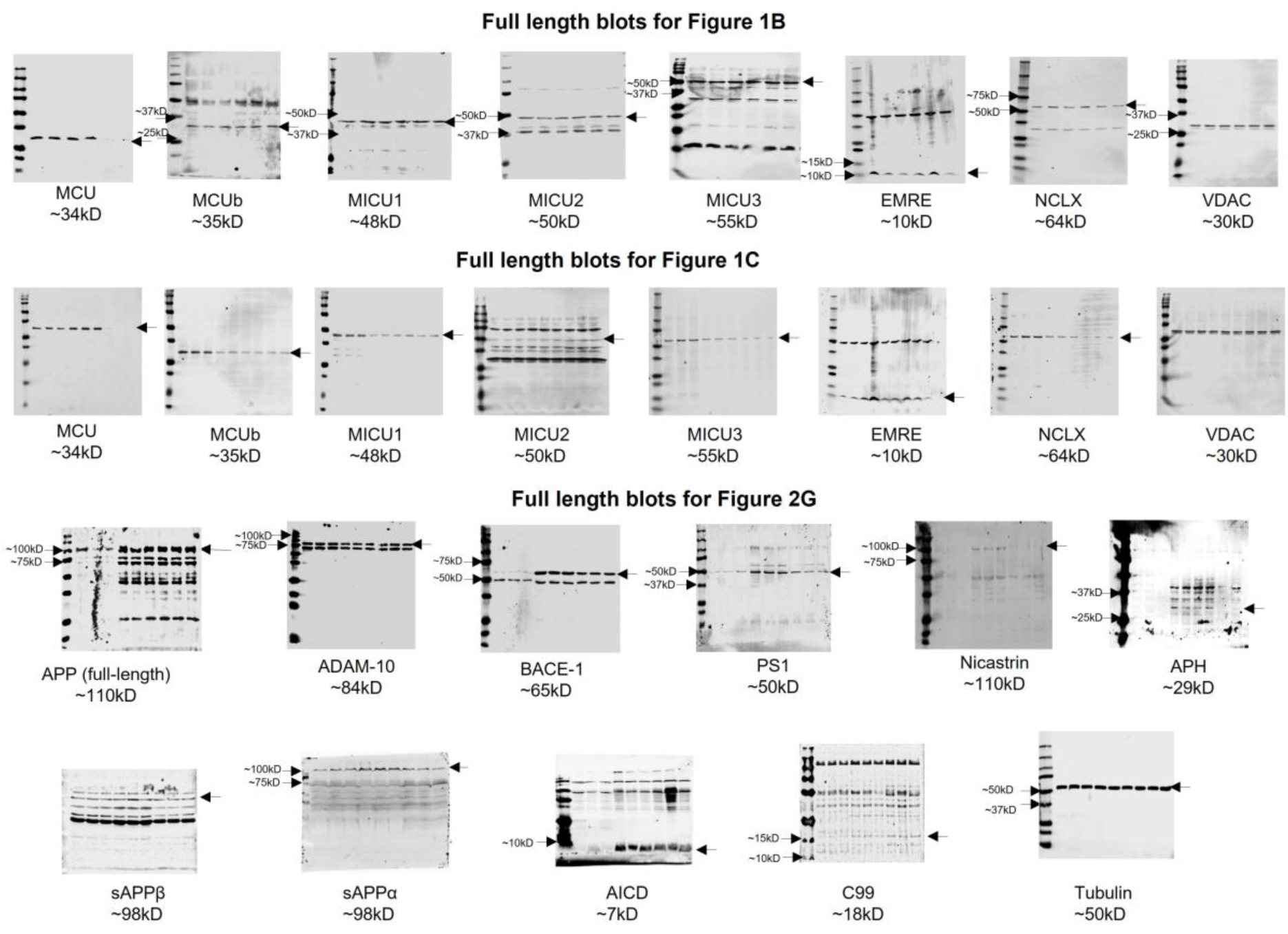
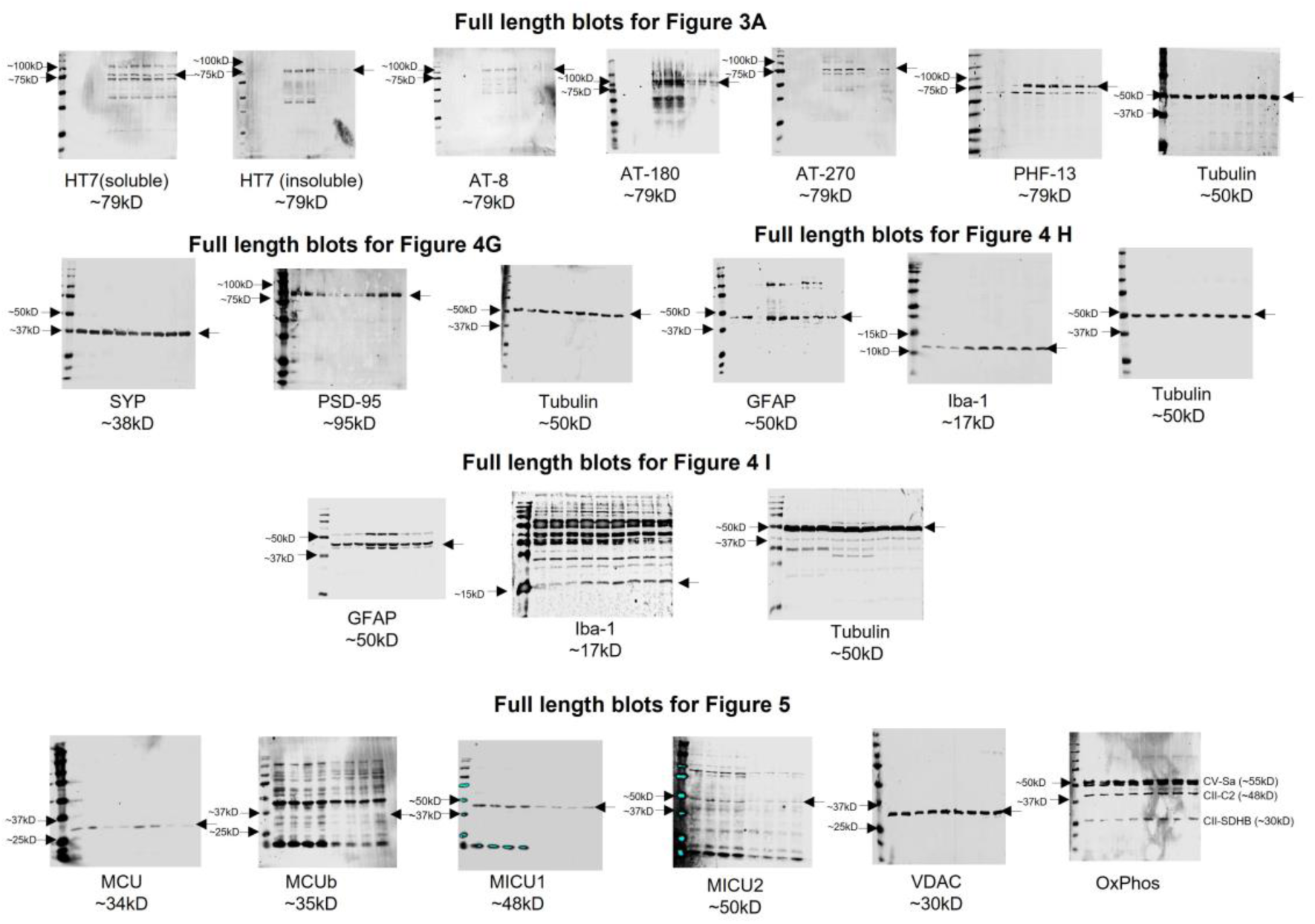
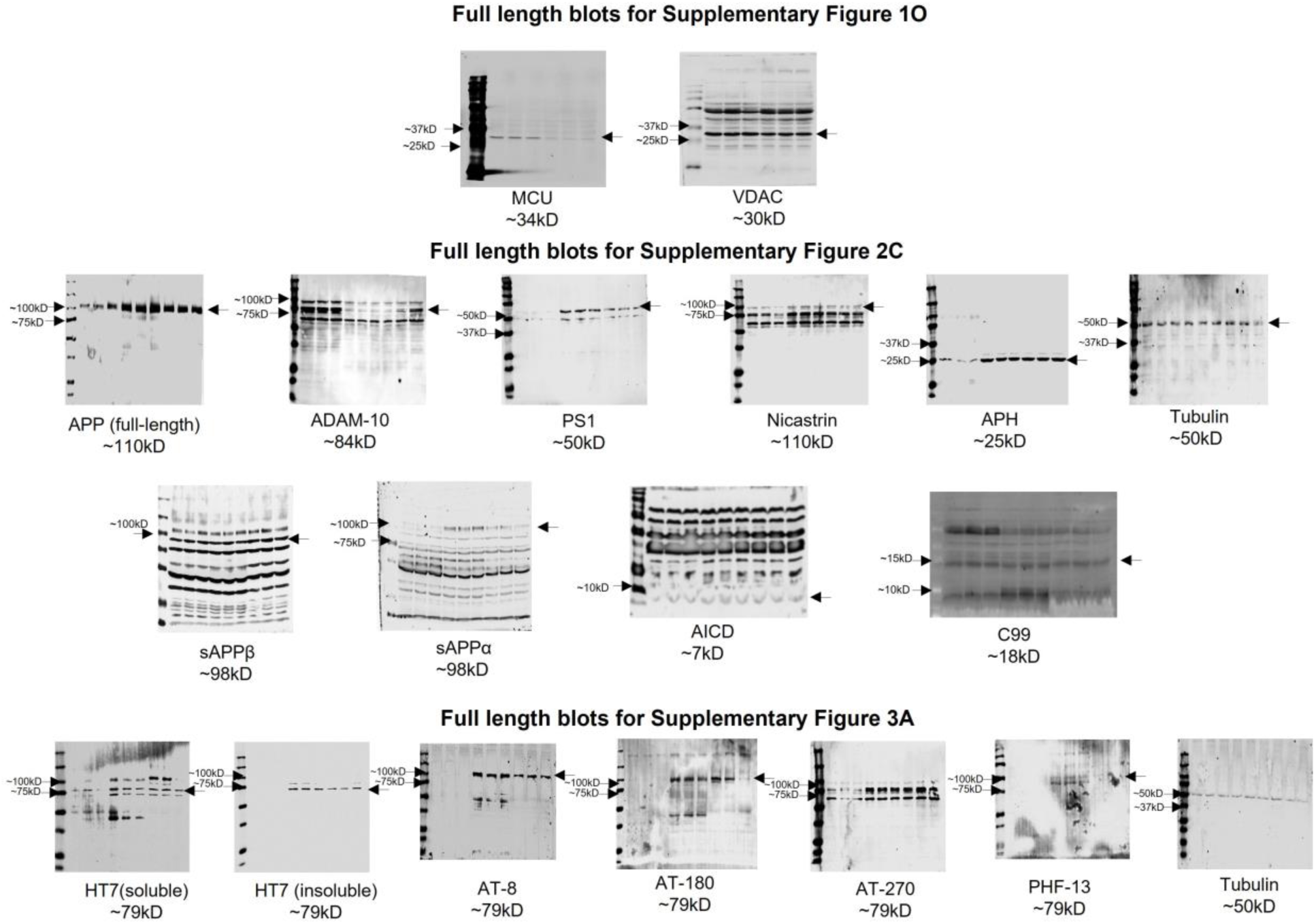
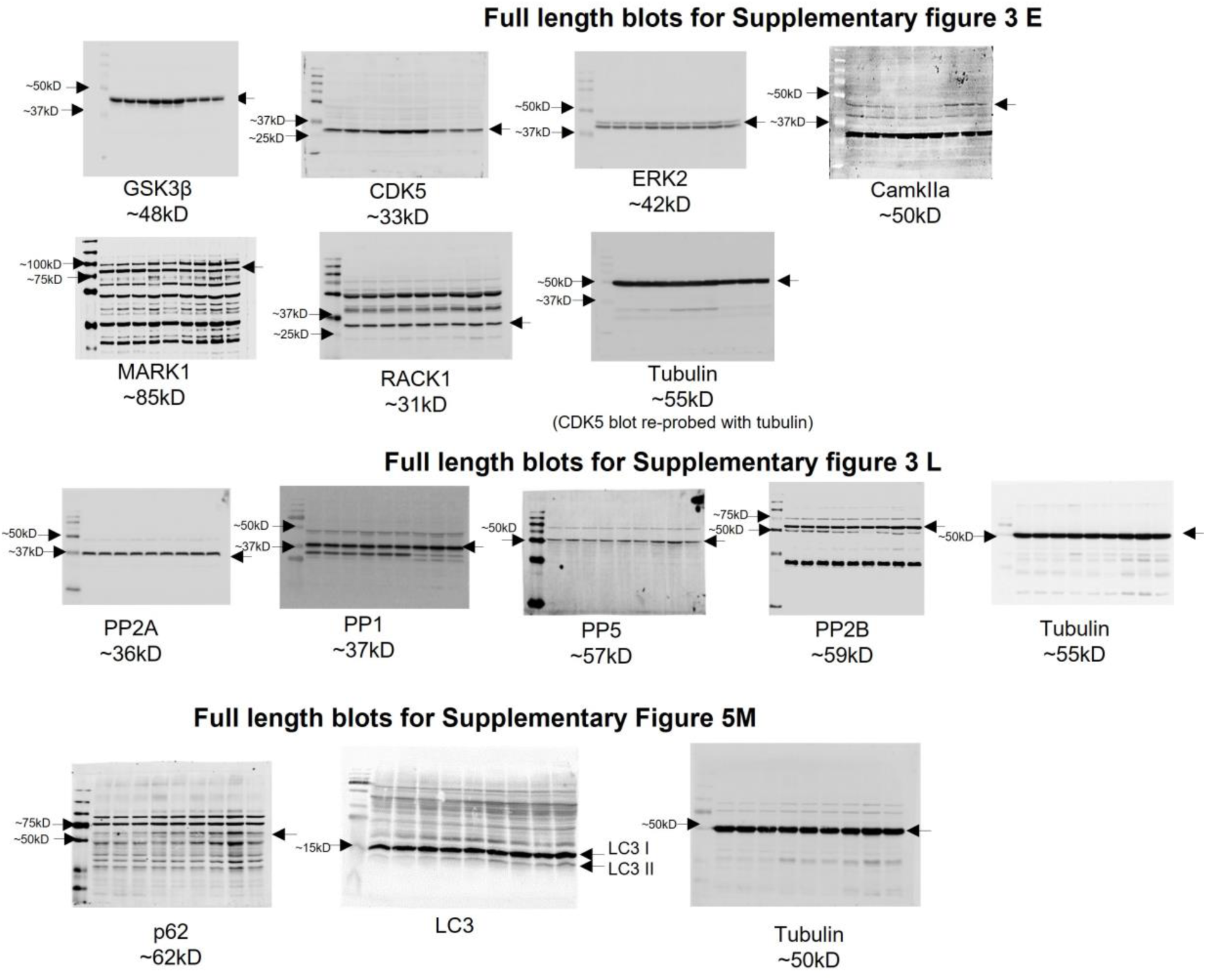
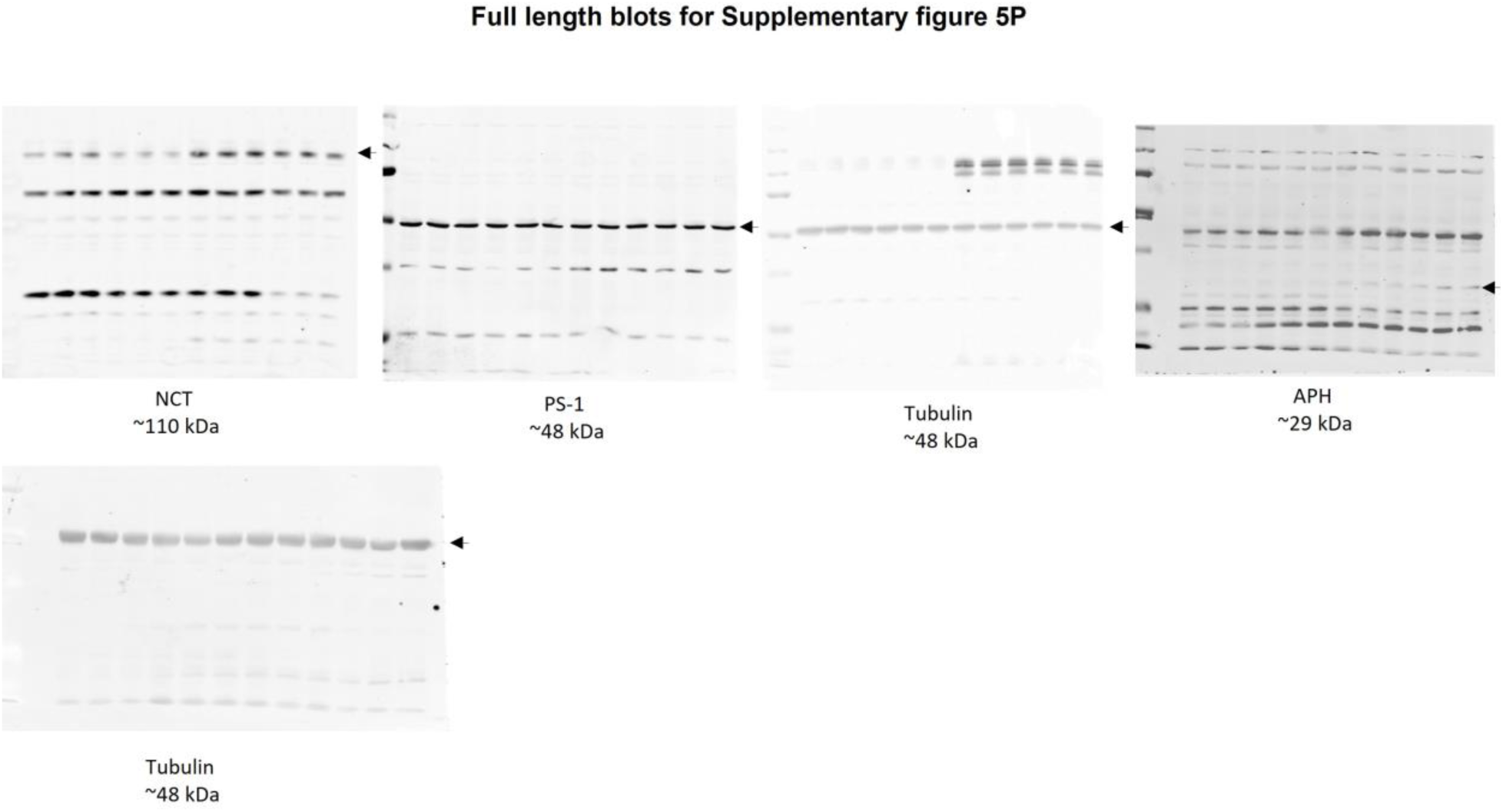
Full-length Western blots

## REFERENCES

1. Almeida CG, Tampellini D, Takahashi RH, Greengard P, Lin MT, Snyder EM, Gouras GK (2005) Beta-amyloid accumulation in APP mutant neurons reduces PSD-95 and GluR1 in synapses. Neurobiology of disease 20: 187–198

2. Area-Gomez E, de Groof A, Bonilla E, Montesinos J, Tanji K, Boldogh I, Pon L, Schon EA (2018) A key role for MAM in mediating mitochondrial dysfunction in Alzheimer disease. Cell Death Dis 9: 335

3. Area-Gomez E, Del Carmen, Lara Castillo M, Tambini MD, Guardia-Laguarta C, de Groof AJ, Madra M, Ikenouchi J, Umeda M, Bird TD, Sturley SL et al (2012) Upregulated function of mitochondria-associated ER membranes in Alzheimer disease. The EMBO journal 31: 4106–4123

4. Ashrafi G, de Juan-Sanz J, Farrell RJ, Ryan TA (2020) Molecular Tuning of the Axonal Mitochondrial Ca(2+) Uniporter Ensures Metabolic Flexibility of Neurotransmission. Neuron 105: 678–687 e675

5. Barrett EF, Barrett JN, David G (2014) Dysfunctional mitochondrial Ca(2+) handling in mutant SOD1 mouse models of fALS: integration of findings from motor neuron somata and motor terminals. Frontiers in cellular neuroscience 8: 184

6. Baughman JM, Perocchi F, Girgis HS, Plovanich M, Belcher-Timme CA, Sancak Y, Bao XR, Strittmatter L, Goldberger O, Bogorad RL et al (2011) Integrative genomics identifies MCU as an essential component of the mitochondrial calcium uniporter. Nature 476: 341–345

7. Begley JG, Duan W, Chan S, Duff K, Mattson MP (1999) Altered calcium homeostasis and mitochondrial dysfunction in cortical synaptic compartments of presenilin-1 mutant mice. J Neurochem 72: 1030–1039

8. Benarroch EE (2018) Glutamatergic synaptic plasticity and dysfunction in Alzheimer disease: Emerging mechanisms. Neurology 91: 125–132

9. Billups B, Forsythe ID (2002) Presynaptic mitochondrial calcium sequestration influences transmission at mammalian central synapses. J Neurosci 22: 5840–5847

10. Butterfield DA, Boyd-Kimball D (2018) Oxidative Stress, Amyloid-beta Peptide, and Altered Key Molecular Pathways in the Pathogenesis and Progression of Alzheimer’s Disease. Journal of Alzheimer’s disease: JAD 62: 1345–1367

11. Cai H QJ, Chen S, Yang J, Hölscher C, Wang Z, Qi J, Wu M. (2022) MCU knockdown in hippocampal neurons improves memory performance of an Alzheimer’s disease mouse model. Acta Biochim Biophys Sin (Shanghai): 1528–1539

12. Call JA, Nichenko AS (2020) Autophagy: an essential but limited cellular process for timely skeletal muscle recovery from injury. Autophagy 16: 1344–1347

13. Calvo-Rodriguez M, Hernando-Perez E, Nunez L, Villalobos C (2019) Amyloid beta Oligomers Increase ER-Mitochondria Ca(2+) Cross Talk in Young Hippocampal Neurons and Exacerbate Aging-Induced Intracellular Ca(2+) Remodeling. Frontiers in cellular neuroscience 13: 22

14. Calvo-Rodriguez M, Hou SS, Snyder AC, Kharitonova EK, Russ AN, Das S, Fan Z, Muzikansky A, Garcia-Alloza M, Serrano-Pozo A et al (2020) Increased mitochondrial calcium levels associated with neuronal death in a mouse model of Alzheimer’s disease. Nat Commun 11: 2146

15. Cardenas C, Miller RA, Smith I, Bui T, Molgo J, Muller M, Vais H, Cheung KH, Yang J, Parker I et al (2010) Essential regulation of cell bioenergetics by constitutive InsP3 receptor Ca2+ transfer to mitochondria. Cell 142: 270–283

16. Caruso D, Barron AM, Brown MA, Abbiati F, Carrero P, Pike CJ, Garcia-Segura LM, Melcangi RC (2013) Age-related changes in neuroactive steroid levels in 3xTg-AD mice. Neurobiology of aging 34: 1080–1089

17. Chang EH, Savage MJ, Flood DG, Thomas JM, Levy RB, Mahadomrongkul V, Shirao T, Aoki C, Huerta PT (2006) AMPA receptor downscaling at the onset of Alzheimer’s disease pathology in double knockin mice. Proc Natl Acad Sci U S A 103: 3410–3415

18. Cheung KH, Mei L, Mak DO, Hayashi I, Iwatsubo T, Kang DE, Foskett JK (2010) Gain-of-function enhancement of IP3 receptor modal gating by familial Alzheimer’s disease-linked presenilin mutants in human cells and mouse neurons. Sci Signal 3: ra22

19. Cheung KH, Shineman D, Muller M, Cardenas C, Mei L, Yang J, Tomita T, Iwatsubo T, Lee VM, Foskett JK (2008) Mechanism of Ca2+ disruption in Alzheimer’s disease by presenilin regulation of InsP3 receptor channel gating. Neuron 58: 871–883

20. Cruz JC, Tseng HC, Goldman JA, Shih H, Tsai LH (2003) Aberrant Cdk5 activation by p25 triggers pathological events leading to neurodegeneration and neurofibrillary tangles. Neuron 40: 471–483

21. Curzon P, Rustay NR, Browman KE (2009) Cued and Contextual Fear Conditioning for Rodents. In: Methods of Behavior Analysis in Neuroscience, nd, Buccafusco J.J. (eds.)Boca Raton (FL)

22. De Stefani D, Raffaello A, Teardo E, Szabo I, Rizzuto R (2011) A forty-kilodalton protein of the inner membrane is the mitochondrial calcium uniporter. Nature 476: 336–340

23. Denton RM, Randle PJ, Martin BR (1972) Stimulation by calcium ions of pyruvate dehydrogenase phosphate phosphatase. Biochem J 128: 161–163

24. Denton RM, Richards DA, Chin JG (1978) Calcium ions and the regulation of NAD+-linked isocitrate dehydrogenase from the mitochondria of rat heart and other tissues. Biochem J 176: 899–906

25. Dreses-Werringloer U, Lambert JC, Vingtdeux V, Zhao H, Vais H, Siebert A, Jain A, Koppel J, Rovelet-Lecrux A, Hannequin D et al (2008) A polymorphism in CALHM1 influences Ca2+ homeostasis, Abeta levels, and Alzheimer’s disease risk. Cell 133: 1149–1161

26. Du H, Guo L, Fang F, Chen D, Sosunov AA, McKhann GM, Yan Y, Wang C, Zhang H, Molkentin JD et al (2008) Cyclophilin D deficiency attenuates mitochondrial and neuronal perturbation and ameliorates learning and memory in Alzheimer’s disease. Nature medicine 14: 1097–1105

27. El-Husseini AE, Schnell E, Chetkovich DM, Nicoll RA, Bredt DS (2000) PSD-95 involvement in maturation of excitatory synapses. Science 290: 1364–1368

28. Evangelopoulos ME, Weis J, Kruttgen A (2005) Signalling pathways leading to neuroblastoma differentiation after serum withdrawal: HDL blocks neuroblastoma differentiation by inhibition of EGFR. Oncogene 24: 3309–3318

29. Fan MM, Raymond LA (2007) N-methyl-D-aspartate (NMDA) receptor function and excitotoxicity in Huntington’s disease. Progress in neurobiology 81: 272–293

30. Ferreira IL, Ferreiro E, Schmidt J, Cardoso JM, Pereira CM, Carvalho AL, Oliveira CR, Rego AC (2015) Abeta and NMDAR activation cause mitochondrial dysfunction involving ER calcium release. Neurobiology of aging 36: 680–692

31. Gabuzda D, Busciglio J, Chen LB, Matsudaira P, Yankner BA (1994) Inhibition of energy metabolism alters the processing of amyloid precursor protein and induces a potentially amyloidogenic derivative. The Journal of biological chemistry 269: 13623–13628

32. Gandhi S, Wood-Kaczmar A, Yao Z, Plun-Favreau H, Deas E, Klupsch K, Downward J, Latchman DS, Tabrizi SJ, Wood NW et al (2009) PINK1-associated Parkinson’s disease is caused by neuronal vulnerability to calcium-induced cell death. Mol Cell 33: 627–638

33. Gillardon F, Rist W, Kussmaul L, Vogel J, Berg M, Danzer K, Kraut N, Hengerer B (2007) Proteomic and functional alterations in brain mitochondria from Tg2576 mice occur before amyloid plaque deposition. Proteomics 7: 605–616

34. Granatiero V, Pacifici M, Raffaello A, De Stefani D, Rizzuto R (2019) Overexpression of Mitochondrial Calcium Uniporter Causes Neuronal Death. Oxid Med Cell Longev 2019: 1681254

35. Gylys KH, Fein JA, Yang F, Wiley DJ, Miller CA, Cole GM (2004) Synaptic changes in Alzheimer’s disease: increased amyloid-beta and gliosis in surviving terminals is accompanied by decreased PSD-95 fluorescence. The American journal of pathology 165: 1809–1817

36. Habib N, McCabe C, Medina S, Varshavsky M, Kitsberg D, Dvir-Szternfeld R, Green G, Dionne D, Nguyen L, Marshall JL et al (2020) Disease-associated astrocytes in Alzheimer’s disease and aging. Nat Neurosci 23: 701–706

37. Harris JJ, Jolivet R, Attwell D (2012) Synaptic energy use and supply. Neuron 75: 762–777

38. Hong S, Beja-Glasser VF, Nfonoyim BM, Frouin A, Li S, Ramakrishnan S, Merry KM, Shi Q, Rosenthal A, Barres BA et al (2016) Complement and microglia mediate early synapse loss in Alzheimer mouse models. Science 352: 712–716

39. Iijima K, Ando K, Takeda S, Satoh Y, Seki T, Itohara S, Greengard P, Kirino Y, Nairn AC, Suzuki T (2000) Neuron-specific phosphorylation of Alzheimer’s beta-amyloid precursor protein by cyclin-dependent kinase 5. J Neurochem 75: 1085–1091

40. Jackson J, Jambrina E, Li J, Marston H, Menzies F, Phillips K, Gilmour G (2019) Targeting the Synapse in Alzheimer’s Disease. Front Neurosci 13: 735

41. Jadiya P, Cohen HM, Kolmetzky DW, Kadam AA, Tomar D, Elrod JW (2023a) Neuronal loss of NCLX-dependent mitochondrial calcium efflux mediates age-associated cognitive decline. iScience 26: 106296

42. Jadiya P, Kolmetzky DW, Tomar D, Di Meco A, Lombardi AA, Lambert JP, Luongo TS, Ludtmann MH, Pratico D, Elrod JW (2019) Impaired mitochondrial calcium efflux contributes to disease progression in models of Alzheimer’s disease. Nat Commun 10: 3885

43. Jadiya P, Kolmetzky DW, Tomar D, Thomas M, Cohen HM, Khaledi S, Garbincius JF, Hildebrand AN, Elrod JW (2023b) Genetic ablation of neuronal mitochondrial calcium uptake halts Alzheimer’s disease progression. bioRxiv

44. Jo DG, Arumugam TV, Woo HN, Park JS, Tang SC, Mughal M, Hyun DH, Park JH, Choi YH, Gwon AR et al (2010) Evidence that gamma-secretase mediates oxidative stress-induced beta-secretase expression in Alzheimer’s disease. Neurobiology of aging 31: 917–925

45. Kim E, Cho KO, Rothschild A, Sheng M (1996) Heteromultimerization and NMDA receptor-clustering activity of Chapsyn-110, a member of the PSD-95 family of proteins. Neuron 17: 103–113

46. Kipanyula MJ, Contreras L, Zampese E, Lazzari C, Wong AK, Pizzo P, Fasolato C, Pozzan T (2012) Ca2+ dysregulation in neurons from transgenic mice expressing mutant presenilin 2. Aging cell 11: 885–893

47. Kostic M, Ludtmann MH, Bading H, Hershfinkel M, Steer E, Chu CT, Abramov AY, Sekler I (2015) PKA Phosphorylation of NCLX Reverses Mitochondrial Calcium Overload and Depolarization, Promoting Survival of PINK1-Deficient Dopaminergic Neurons. Cell Rep 13: 376–386

48. Lacampagne A, Liu X, Reiken S, Bussiere R, Meli AC, Lauritzen I, Teich AF, Zalk R, Saint N, Arancio O et al (2017) Post-translational remodeling of ryanodine receptor induces calcium leak leading to Alzheimer’s disease-like pathologies and cognitive deficits. Acta Neuropathol 134: 749–767

49. Lau A, Tymianski M (2010) Glutamate receptors, neurotoxicity and neurodegeneration. Pflugers Archiv: European journal of physiology 460: 525–542

50. Liao Y, Hao Y, Chen H, He Q, Yuan Z, Cheng J (2015) Mitochondrial calcium uniporter protein MCU is involved in oxidative stress-induced cell death. Protein & cell 6: 434–442

51. Llorens-Martin M, Fuster-Matanzo A, Teixeira CM, Jurado-Arjona J, Ulloa F, Defelipe J, Rabano A, Hernandez F, Soriano E, Avila J (2013) Alzheimer disease-like cellular phenotype of newborn granule neurons can be reversed in GSK-3beta-overexpressing mice. Mol Psychiatry 18: 395

52. Logan CV, Szabadkai G, Sharpe JA, Parry DA, Torelli S, Childs AM, Kriek M, Phadke R, Johnson CA, Roberts NY et al (2014) Loss-of-function mutations in MICU1 cause a brain and muscle disorder linked to primary alterations in mitochondrial calcium signaling. Nature genetics 46: 188–193

53. Luongo TS, Lambert JP, Gross P, Nwokedi M, Lombardi AA, Shanmughapriya S, Carpenter AC, Kolmetzky D, Gao E, van Berlo JH et al (2017) The mitochondrial Na(+)/Ca(2+) exchanger is essential for Ca(2+) homeostasis and viability. Nature 545: 93–97

54. Luongo TS, Lambert JP, Yuan A, Zhang X, Gross P, Song J, Shanmughapriya S, Gao E, Jain M, Houser SR et al (2015) The Mitochondrial Calcium Uniporter Matches Energetic Supply with Cardiac Workload during Stress and Modulates Permeability Transition. Cell Rep 12: 23–34

55. Lustbader JW, Cirilli M, Lin C, Xu HW, Takuma K, Wang N, Caspersen C, Chen X, Pollak S, Chaney M et al (2004) ABAD directly links Abeta to mitochondrial toxicity in Alzheimer’s disease. Science 304: 448–452

56. Mallilankaraman K, Doonan P, Cardenas C, Chandramoorthy HC, Muller M, Miller R, Hoffman NE, Gandhirajan RK, Molgo J, Birnbaum MJ et al (2012) MICU1 is an essential gatekeeper for MCU-mediated mitochondrial Ca(2+) uptake that regulates cell survival. Cell 151: 630–644

57. Mattson MP, Cheng B, Davis D, Bryant K, Lieberburg I, Rydel RE (1992) beta-Amyloid peptides destabilize calcium homeostasis and render human cortical neurons vulnerable to excitotoxicity. J Neurosci 12: 376–389

58. Migaud M, Charlesworth P, Dempster M, Webster LC, Watabe AM, Makhinson M, He Y, Ramsay MF, Morris RG, Morrison JH et al (1998) Enhanced long-term potentiation and impaired learning in mice with mutant postsynaptic density-95 protein. Nature 396: 433–439

59. Muller M, Cheung KH, Foskett JK (2011) Enhanced ROS generation mediated by Alzheimer’s disease presenilin regulation of InsP3R Ca2+ signaling. Antioxidants & redox signaling 14: 1225–1235

60. Nakagawa T, Zhu H, Morishima N, Li E, Xu J, Yankner BA, Yuan J (2000) Caspase-12 mediates endoplasmic-reticulum-specific apoptosis and cytotoxicity by amyloid-beta. Nature 403: 98–103

61. Nichenko AS, Southern WM, Tehrani KF, Qualls AE, Flemington AB, Mercer GH, Yin A, Mortensen LJ, Yin H, Call JA (2020) Mitochondrial-specific autophagy linked to mitochondrial dysfunction following traumatic freeze injury in mice. Am J Physiol Cell Physiol 318: C242–C252

62. Oddo S, Caccamo A, Shepherd JD, Murphy MP, Golde TE, Kayed R, Metherate R, Mattson MP, Akbari Y, LaFerla FM (2003) Triple-transgenic model of Alzheimer’s disease with plaques and tangles: intracellular Abeta and synaptic dysfunction. Neuron 39: 409–421

63. Orr AL, Kim C, Jimenez-Morales D, Newton BW, Johnson JR, Krogan NJ, Swaney DL, Mahley RW (2019) Neuronal Apolipoprotein E4 Expression Results in Proteome-Wide Alterations and Compromises Bioenergetic Capacity by Disrupting Mitochondrial Function. Journal of Alzheimer’s disease: JAD 68: 991–1011

64. Oyarzabal A, Marin-Valencia I (2019) Synaptic energy metabolism and neuronal excitability, in sickness and health. J Inherit Metab Dis 42: 220–236

65. Panel M, Ghaleh B, Morin D (2018) Mitochondria and aging: A role for the mitochondrial transition pore? Aging cell 17: e12793

66. Park JS, Kam TI, Lee S, Park H, Oh Y, Kwon SH, Song JJ, Kim D, Kim H, Jhaldiyal A et al (2021) Blocking microglial activation of reactive astrocytes is neuroprotective in models of Alzheimer’s disease. Acta Neuropathol Commun 9: 78

67. Paula-Lima AC, Adasme T, SanMartin C, Sebollela A, Hetz C, Carrasco MA, Ferreira ST, Hidalgo C (2011) Amyloid beta-peptide oligomers stimulate RyR-mediated Ca2+ release inducing mitochondrial fragmentation in hippocampal neurons and prevent RyR-mediated dendritic spine remodeling produced by BDNF. Antioxidants & redox signaling 14: 1209–1223

68. Perez MJ, Ponce DP, Aranguiz A, Behrens MI, Quintanilla RA (2018) Mitochondrial permeability transition pore contributes to mitochondrial dysfunction in fibroblasts of patients with sporadic Alzheimer’s disease. Redox Biol 19: 290–300

69. Perocchi F, Gohil VM, Girgis HS, Bao XR, McCombs JE, Palmer AE, Mootha VK (2010) MICU1 encodes a mitochondrial EF hand protein required for Ca(2+) uptake. Nature 467: 291–296

70. Petersen A, Castilho RF, Hansson O, Wieloch T, Brundin P (2000) Oxidative stress, mitochondrial permeability transition and activation of caspases in calcium ionophore A23187-induced death of cultured striatal neurons. Brain research 857: 20–29

71. Plovanich M, Bogorad RL, Sancak Y, Kamer KJ, Strittmatter L, Li AA, Girgis HS, Kuchimanchi S, De Groot J, Speciner L et al (2013) MICU2, a paralog of MICU1, resides within the mitochondrial uniporter complex to regulate calcium handling. PLoS One 8: e55785

72. Qiu J, Tan YW, Hagenston AM, Martel MA, Kneisel N, Skehel PA, Wyllie DJ, Bading H, Hardingham GE (2013) Mitochondrial calcium uniporter Mcu controls excitotoxicity and is transcriptionally repressed by neuroprotective nuclear calcium signals. Nat Commun 4: 2034

73. Raffaello A, De Stefani D, Sabbadin D, Teardo E, Merli G, Picard A, Checchetto V, Moro S, Szabo I, Rizzuto R (2013) The mitochondrial calcium uniporter is a multimer that can include a dominant-negative pore-forming subunit. The EMBO journal 32: 2362–2376

74. Rhein V, Song X, Wiesner A, Ittner LM, Baysang G, Meier F, Ozmen L, Bluethmann H, Drose S, Brandt U et al (2009) Amyloid-beta and tau synergistically impair the oxidative phosphorylation system in triple transgenic Alzheimer’s disease mice. Proc Natl Acad Sci U S A 106: 20057–20062

75. Rossi A, Rigotto G, Valente G, Giorgio V, Basso E, Filadi R, Pizzo P (2020) Defective Mitochondrial Pyruvate Flux Affects Cell Bioenergetics in Alzheimer’s Disease-Related Models. Cell Rep 30: 2332–2348 e2310

76. Sarasija S, Laboy JT, Ashkavand Z, Bonner J, Tang Y, Norman KR (2018) Presenilin mutations deregulate mitochondrial Ca(2+) homeostasis and metabolic activity causing neurodegeneration in Caenorhabditis elegans. eLife 7

77. Singh R, Bartok A, Paillard M, Tyburski A, Elliott M, Hajnoczky G (2022) Uncontrolled mitochondrial calcium uptake underlies the pathogenesis of neurodegeneration in MICU1 -deficient mice and patients. Sci Adv 8: eabj4716

78. Soman S, Keatinge M, Moein M, Da Costa M, Mortiboys H, Skupin A, Sugunan S, Bazala M, Kuznicki J, Bandmann O (2017) Inhibition of the mitochondrial calcium uniporter rescues dopaminergic neurons in pink1(-/-) zebrafish. Eur J Neurosci 45: 528–535

79. Soman SK, Bazala M, Keatinge M, Bandmann O, Kuznicki J (2019) Restriction of mitochondrial calcium overload by mcu inactivation renders a neuroprotective effect in zebrafish models of Parkinson’s disease. Biol Open 8

80. Starkov AA, Chinopoulos C, Fiskum G (2004) Mitochondrial calcium and oxidative stress as mediators of ischemic brain injury. Cell calcium 36: 257–264

81. Stout AK, Raphael HM, Kanterewicz BI, Klann E, Reynolds IJ (1998) Glutamate-induced neuron death requires mitochondrial calcium uptake. Nat Neurosci 1: 366–373

82. Stutzmann GE, Caccamo A, LaFerla FM, Parker I (2004) Dysregulated IP3 signaling in cortical neurons of knock-in mice expressing an Alzheimer’s-linked mutation in presenilin1 results in exaggerated Ca2+ signals and altered membrane excitability. J Neurosci 24: 508–513

83. Szalai G, Krishnamurthy R, Hajnoczky G (1999) Apoptosis driven by IP(3)-linked mitochondrial calcium signals. The EMBO journal 18: 6349–6361

84. Tarsa L, Goda Y (2002) Synaptophysin regulates activity-dependent synapse formation in cultured hippocampal neurons. Proc Natl Acad Sci U S A 99: 1012–1016

85. Thinakaran G, Teplow DB, Siman R, Greenberg B, Sisodia SS (1996) Metabolism of the “Swedish” amyloid precursor protein variant in neuro2a (N2a) cells. Evidence that cleavage at the “beta-secretase” site occurs in the golgi apparatus. The Journal of biological chemistry 271: 9390–9397

86. Tong BC, Lee CS, Cheng WH, Lai KO, Foskett JK, Cheung KH (2016) Familial Alzheimer’s disease-associated presenilin 1 mutants promote gamma-secretase cleavage of STIM1 to impair store-operated Ca2+ entry. Sci Signal 9: ra89

87. Tsien JZ, Chen DF, Gerber D, Tom C, Mercer EH, Anderson DJ, Mayford M, Kandel ER, Tonegawa S (1996) Subregion-and cell type-restricted gene knockout in mouse brain. Cell 87: 1317–1326

88. Vergara RC, Jaramillo-Riveri S, Luarte A, Moenne-Loccoz C, Fuentes R, Couve A, Maldonado PE (2019) The Energy Homeostasis Principle: Neuronal Energy Regulation Drives Local Network Dynamics Generating Behavior. Front Comput Neurosci 13: 49

89. Verma M, Callio J, Otero PA, Sekler I, Wills ZP, Chu CT (2017) Mitochondrial Calcium Dysregulation Contributes to Dendrite Degeneration Mediated by PD/LBD-Associated LRRK2 Mutants. J Neurosci 37: 11151–11165

90. Vicente Roca-Agujetas CdD, Laura Lestón, Montserrat Marí, Albert Morales, Anna Colell (2019) Recent Insights into the Mitochondrial Role in Autophagy and Its Regulation by Oxidative Stress. Oxidative Medicine and Cellular Longevity: 16

91. Wu HY, Tomizawa K, Oda Y, Wei FY, Lu YF, Matsushita M, Li ST, Moriwaki A, Matsui H (2004) Critical role of calpain-mediated cleavage of calcineurin in excitotoxic neurodegeneration. The Journal of biological chemistry 279: 4929–4940

92. Xie N, Wu C, Wang C, Cheng X, Zhang L, Zhang H, Lian Y (2017) Inhibition of the mitochondrial calcium uniporter inhibits Abeta-induced apoptosis by reducing reactive oxygen species-mediated endoplasmic reticulum stress in cultured microglia. Brain research 1676: 100–106

